# Comparative study on the inhibition of copper oxide, nickel, and sodium tungstate on microbially induced concrete corrosion under sewer conditions

**DOI:** 10.64898/2026.03.05.709923

**Authors:** Kaijie Wang, Xiaoyan Sun, Kairong Lin, Xiaohong Chen, Qilin Wang

**Affiliations:** School of Civil Engineering, Sun Yat-sen University, Zhuhai, 519082, PR China; Center for Water Resources and Environment Research, Sun Yat-sen University, Guangzhou, 510275, PR China; Centre for Technology in Water and Wastewater, School of Civil and Environmental Engineering, University of Technology Sydney, Ultimo, NSW 2007, Australia

**Keywords:** microbially induced concrete corrosion, sewer systems, metal bacteriostatic agent, copper oxide, nickel, sodium tungstate

## Abstract

Microbially induced concrete corrosion (MICC) is a significant issue that reduces the service life of sewer systems. Bacteriostatic agent in concrete can inhibit microbial activity and the process of MICC to some extent. However, a systematic comparison of the inhibition effects of various bacteriostatic agents on MICC remains lacking. In this study, three bacteriostatic agents (copper oxide, nickel, and sodium tungstate) were investigated for their inhibitory effects on MICC. For each inhibitor, the cement mortar coupons with 0.05 wt%, 0.1 wt%, and 0.2 wt% of the inhibitor were prepared. The coupons were partially submerged in sewage of a controlled laboratory corrosion chamber (20 ± 5 ppm H_2_S) to simulate the tidal region of gravity sewer. During the 56 days of exposure, the intensification of pores, cracks, surface erosion, and spalling was observed on all coupons. After 56 days of exposure, the sulfate concentration and adenosine triphosphate (ATP) content of coupons without inhibitor were 10.65 mg/cm^2^ and 30.17 ± 3.87 μmol/cm^2^, respectively. They were higher than those of coupons containing 0.05 wt%, 0.1 wt%, and 0.2 wt% of copper oxide and 0.05wt% of nickel. The temporal profiles of ATP of coupons without inhibitor was similar to those of coupons containing sodium tungstate. After exposure for 28 days, the surface pH of coupons without inhibitor was 7.45, meanwhile of those coupons containing 0.2 wt% of copper oxide and 0.05 wt% of nickel were 9.42 and 9.93, respectively. Those results indicated that the bacteriostatic effect of copper oxide and nickel (0.05 wt %) was found to be the most prominent. The findings indicate that a single bacteriostatic agent is only effective during specific corrosion stages, suggesting that a combination of multiple agents may be a promising strategy to combat the multi-stage MICC process over the long term. This study provides a theoretical basis for the selection and development of protective materials against concrete corrosion in sewer networks.

## 1. Introduction

Sewer pipes are critical infrastructure for sewage collection and transportation in urbane society [1]. Concrete sewer pipes corrode to varying degrees throughout their lifespan and may experience severe corrosion only after a few years of usage [2]. Corrosion can result in structural collapse of sewer pipes, and hence exfiltration of sewage, escape of volatile compounds to the surrounding environment, and posing a significant threat to public health and the safety of urban areas [3–5]. In Germany, microbial corrosion of building materials accounts for 10%-20% of damage [18]. In Flanders, Belgium, the costs of measures taken to deal with microbial corrosion make up 10% of total sewage treatment costs [19]. In the United States, approximately 50 billion US dollars is spent annually to maintain sewer systems, and most of these funds are spent on direct responses to microbial corrosion [20].Therefore, it is necessary to develop effective measures to prevent or inhibit concrete sewer corrosion.

Metabolic products, such as hydrogen sulfide (H_2_S), biologically produced sulfuric acid, and organic acids, contribute to the corrosion of concrete. Sulfate reducing bacteria (SRB) reduces sulfate in sewage to H_2_S in anaerobic conditions [6, 7]. The transfer of H_2_S and CO_2_ from sewer atmosphere to concrete surface reduces the pH of the concrete surface from 13-11 to approximately 9 [8]. When the pH of the concrete surface is between 9-3, neutrophilic sulfur-oxidizing bacteria (NSOB) colonize the concrete surface and oxidize H_2_S to sulfuric acid [9, 10]. This further reduces surface pH and results in the colonization of acidophilic sulfur-oxidizing bacteria (ASOB) [11, 12]. Sulfuric acid reacts with concrete to form gypsum, ettringite, and other products. These products cause volume expansion and structural damage to concrete [13]. Additionally, nitric acid is produced by nitrifying bacteria during the nitrification of amine, causing further corrosion to concrete [14–17].

To mitigate microbially induced concrete corrosion (MICC), various bacteriostatic agents, such as organic and inorganic compounds, have been utilized to inhibit the growth and activity of microbes [21]. Organic compounds could be unstable under temperature changes due to thermal degradation. For a long-term application, their antibacterial effect may decrease due to changes in environmental factors [22]. Furthermore, microorganisms can develop resistance to organic bacteriostatic agents, causing bacteriostatic agents to lose their bacteriostatic effect. Inorganic compounds, e.g. metals and metallic compounds, have a long period of validity and are not easy to fail due to temperature changes. But they may have a certain level of toxicity and can pollute the environment [23–28]. Inorganic bacteriostatic agents can be more effective at inhibiting bacterial growth than organic bacteriostatic agents, provided that their dosage is carefully controlled within safe environmental limits [9, 29].

Research by Alum et al. [30] demonstrated that adding 20% zinc oxide to concrete was the most effective in inhibiting algae growth compared with the coupons containing sodium bromide (SB), ammonium chloride (AC), copper slag (CS), and cetyl-methyl-ammonium bromide (CAB). Delgado et al. [31] observed that the polypropylene matrix containing either nano-copper or nano-copper oxide had antibacterial properties. In addition, the antibacterial effect of copper oxide is better than that of copper, probably due to that ion release rate of the former is higher than that of the latter. Negishi et al. [32] found that adding 0.075% calcium tungstate or 0.075% nickel metal to concrete significantly mitigates MICC during a two-year exposure experiment in real sewers.

Many studies confirmed that calcium tungstate, nickel, copper oxide, and other metallic compounds have inhibitial and/or antibacterial effects on microbial activity. However, the dosage of these metals varies widely among different studies ranging from 0.05 to 30% [25, 30, 32–38]. While higher dosages may yield more significant MICC inhibition, the potential negative environmental impact must be considered. Few studies have explored optimal dosages comprehensively. Therefore, optimizing the dosage of experimentally validated bacteriostatic agents is necessary to achieve the most effective MICC inhibition while minimizing environmental concerns.

This study aims at evaluating the inhibiting impact of copper oxide, nickel, and sodium tungstate at different content on the MICC. These agents were added to cement mortar at cement mass fractions of 0.05%, 0.1%, and 0.2%, respectively. The cement mortar coupon samples were partially submerged in a laboratory corrosion chamber, with the environmental parameters controlled at 20 ± 2L, 90-100% relative humidity, and 20 ± 5 ppm H_2_S. The side surface of the coupons simulated tidal region of a gravity sewer environment. The corrosion of the coupons was monitored of the physical, chemical and microbial activities (e.g. surface pH, sulfate content, mass change, adenosine triphosphate content, etc.) [39, 40].

## 2. Methods and Materials

### 2.1 Cement mortar coupon

Cement mortar coupons with dimension of 4cmc × 4cm × 4cm were prepared. The composition of the mortar coupons and the addition of metal compounds are shown in Table 1. Specifically, the ratio of cement-to-sand-to-water is 2:5:1. China’s ISO standard sand (Standard sand, Xiamen Aisiou) and Portland cement 42.5 (PO42.5 cement, Jiangmen Hailuo) were used in the preparation. Milli-Q water instead of tap water was used to avoid potential impact of chloride ion on cement mortar. In manufacturing the coupons, precise measurements were achieved by using an electronic balance with an accuracy of 0.01g.

**Table 1.** Conformation of cement mortar coupons (g).

| Coupon<br>groups | Quartz<br>sand | PO42.5<br>Cement | Milli-Q<br>water | Copper<br>oxide<br>powder | Atomization<br>nickel<br>powder | Sodium<br>tungstate<br>powder |
| --- | --- | --- | --- | --- | --- | --- |
| K* | 1350 | 540 | 270 | 0 | 0 | 0 |
| C1* | 1350 | 540 | 270 | 0.27 | 0 | 0 |
| C2* | 1350 | 540 | 270 | 0.54 | 0 | 0 |
| C3* | 1350 | 540 | 270 | 1.08 | 0 | 0 |
| N1* | 1350 | 540 | 270 | 0 | 0.27 | 0 |
| N2* | 1350 | 540 | 270 | 0 | 0.54 | 0 |
| N3* | 1350 | 540 | 270 | 0 | 1.08 | 0 |
| W1* | 1350 | 540 | 270 | 0 | 0 | 0.27 |
| W2* | 1350 | 540 | 270 | 0 | 0 | 0.54 |
| W3* | 1350 | 540 | 270 | 0 | 0 | 1.08 |

\*K: blank control coupons, C1-3: coupons containing 0.05 wt%, 0.1 wt% and 0.2% of copper oxide, N1-3: coupons containing 0.05%, 0.1% and 0.2% of nickel, W: coupons
containing 0.05%, 0.1% and 0.2% of sodium tungstate.

The physical indexes of the metallic bacteriostatic agents are shown in Table S1 (See Supplementary Information). To ensure the uniform distribution, the weighed sodium tungstate powder was dissolved in 270mL of Milli-Q water. For water-insoluble copper oxide powders and nickel powders, a cement paste mixer (NJ-160A, Wuxi Construction) was used to ensure even distribution in the cement powder. The mixed mortar was then poured into 4cm*4cm*4cm plastic molds and cured in humid air for 24 hours before demolding. Then the coupons were cured in the curing room (ambient 20 ± 2L, 90-100% relative humidity) for 7 days. Finally, 10 groups of coupons were prepared, i.e. coupons as the blank control (Group K), coupons with 0.05%, 0.1% and 0.2% of copper oxide powder (Group C, including C1, C2, and C3), coupons incorporating 0.05%, 0.1% and 0.2% of sodium tungstate powder (Group N, including N1, N2, and N3), and coupons incorporating 0.05%, 0.1% and 0.2% of sodium tungstate powder (Group W, including W1, W2, and W3).

### 2.2 Corrosion chamber and exposure environment

The coupons were exposed in a laboratory corrosion chamber containing 5.0L of domestic sewage. The sewage was collected from a sewer pumping station (Tangjia Town, Zhuhai City, China) and was replaced every two weeks. The relevant characteristics of sewage is shown in Table S2 (See Supplementary Information). The coupons were partially submerged in the sewage of to a depth of 2 cm. This arrangement simulates the environment of the gravity sewer pipe.

Hydrogen sulfide gas was generated in the reactor by adding 1mol/L sodium sulfide solution to react with excess hydrochloric acid. To maintain the concentration of hydrogen sulfide gas at 20 ± 5 ppm, a programmable logic controller (PLC) was used to control the peristaltic pump (BT100 - 1F, Longer), which dosed the sodium sulfide solution. A hydrogen sulfide gas detector (Man - PPM Rev 6, Acrulog) was used to monitor the gaseous hydrogen sulfide concentration in the reactor. Environmental conditions were checked daily to ensure the proper operation of the corrosion reactor.

### 2.3 Analysis procedure

Every 14 days, ten groups of coupons (K, C1, C2, C3, N1, N2, N3, W1, W2, W3) were retrieved from the reactor for analysis of the following parameters to monitor corrosion activity.

#### 2.3.1 Surface Appearance Changes

Before exposure to the laboratory corrosion chamber and after each sampling event, the coupons were placed on white sterile paper, and their top and side surfaces were photographed. This allowed for monitoring of the temporal and spatial changes in corrosion activity.

#### 2.3.2 Surface pH

The surface pH was measured on both the top and side surface of each coupon. For each surface, three random points were selected, and the final result was determined by averaging these three measurements. Prior to each measurement, the randomly selected spot of the coupon surface was moistened with 0.5 mL of deionized water. After 10 minutes of stabilization, the surface pH was measured by a flat surface pH electrode (E-201-P, Lei-ci).

#### 2.3.3 Sulfate content

The coupon layer was scraped from the coupon surface using a clean scalpel, and the remaining coupon was rinsed with Milli-Q water. The scraped corrosion layer was dissolved in 150mL of Milli-Q water to prepare an eluent. The eluent was thorougly mixed using a vortex to ensure complete solubilization of sulfate from the corrosion layer. Subsequently, the eluent was centrifuged (5810R, Eppendorf) at 1500 rpm for 15 min to separate insoluble solid particles from the liquid phase. The supernatant was collected and then filtered through 0.45 μm filter membranes (622110, SORFA). To prevent potential interference from phenolic compounds, metals and other hydrophobic substances in the supernatant, solid phase extraction cartridges (RP cartridges and Na cartridges, QY3-43001& QY3-46001, QINGYUN) were used for water sample pretreatment. The pretreated supernatant was then analyzed using ionic chromatography (Dionex Aquion, Thermo Scientific™), and the sulfate content in the corrosion layer was determined by quantifying the peak area in the chromatogram.

#### 2.3.4 Mass changes

To quantify the mass change of coupons due to corrosion, the corrosion layer was first scraped off using a scalpel. The coupons were then rinsed with Milli-Q water, and dried in an oven at 60L for 3 days to remove moisture. Then, the mass of each coupon was measured using an electronic balance with an accuracy of 0.01g. The mass change ratio, *R*, was calculated using the following equation:

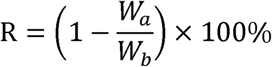

Where *R* is the mass change (%); *W_a_* is the mass of the coupon after exposure (g); *W_b_* is the mass of the coupon before exposure (g).

#### 2.3.5 Characterization of the corrosion layer by Scanning Electron Microscopy and Energy Dispersive X-ray Spectroscopy

To determine the mineral composition of the corrosion layer, Scanning Electron Microscopy (SEM) and Energy Dispersive X-ray Spectroscopy (EDS) were performed on the coupons. Utilizing a gemstone cutting machine (AH1325M, Gxulaser), a longitudinal cut was made along the central region of the coupon. This process yielded a thin section measuring 40 mm*40 mm*3 mm (See Figure S1 in the Supplementary Information). As SEM analysis requires conductive samples, the surface of each section was coated with a layer of carbon. The thin sections were then imaged using the backscattered electron imaging mode of the SEM (SU5000, HITACHI).

EDS analysis was then conducted on the entire region of each SEM image, allowing the determination of elemental composition based on the intensity of the characteristic X-ray signals in the EDS spectrum.

#### 2.3.6 Adenosine triphosphate content

The Adenosine triphosphate (ATP) content of the corrosion layer was determined using a bioluminescence assay. An aliquiot of the corrosion layer eluate (prepared as described in Section 2.3.3) was taken and divided into two portions. One portion was sterilized in an autoclave (IMJ-78A, STIK) at 121 L for 30 minutes to serve as a control, and the other portion was used as the test sample. For analysis, 100 μL of each sample portion was mixed with 100 μL of BacTiter-Glo^TM^ detection reagent (G8231, Promega Corporation) in a 96-well plate (Lumitrac 200, Greiner Bio-one). The plate was then incubated in a microplate reader (Synergy Neo2, BioTek) for 5 minutes, after which the fluorescence value was measured All tests were performed in triplicate.

## 3. Results

### 3.1 Surface morphological changes

Prior to exposure in the corrosion chamber, the top surface of all coupons were similar in color and surface roughness (See Figture S2 in the Supplementary Information). After 7 days of exposure, significant water accumulation was observed on the surface of Group K coupons, resulting in increased localized humidity and higher surface water content (red triangle box in Fig. 1A). Notably, only half of the surface area of coupon K1 exhibited elevated moisture levels, primarily due to its uneven surface topography, leading to greater water accumulation in the lower-lying areas. Subsequent photos confirmed that water evaporated more slowly from these areas, resulting in prolonged water retention.

**Fig. 1.**
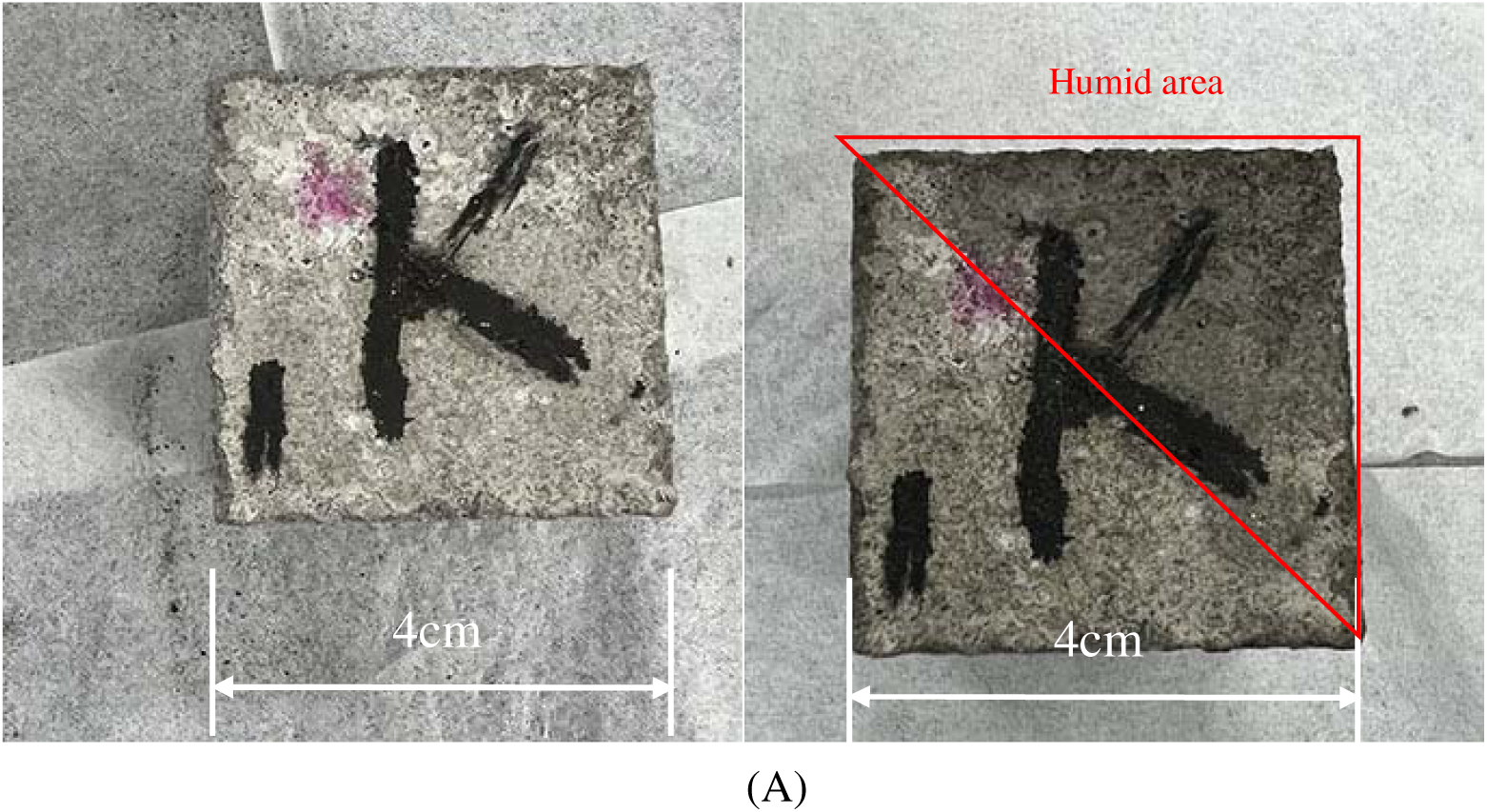
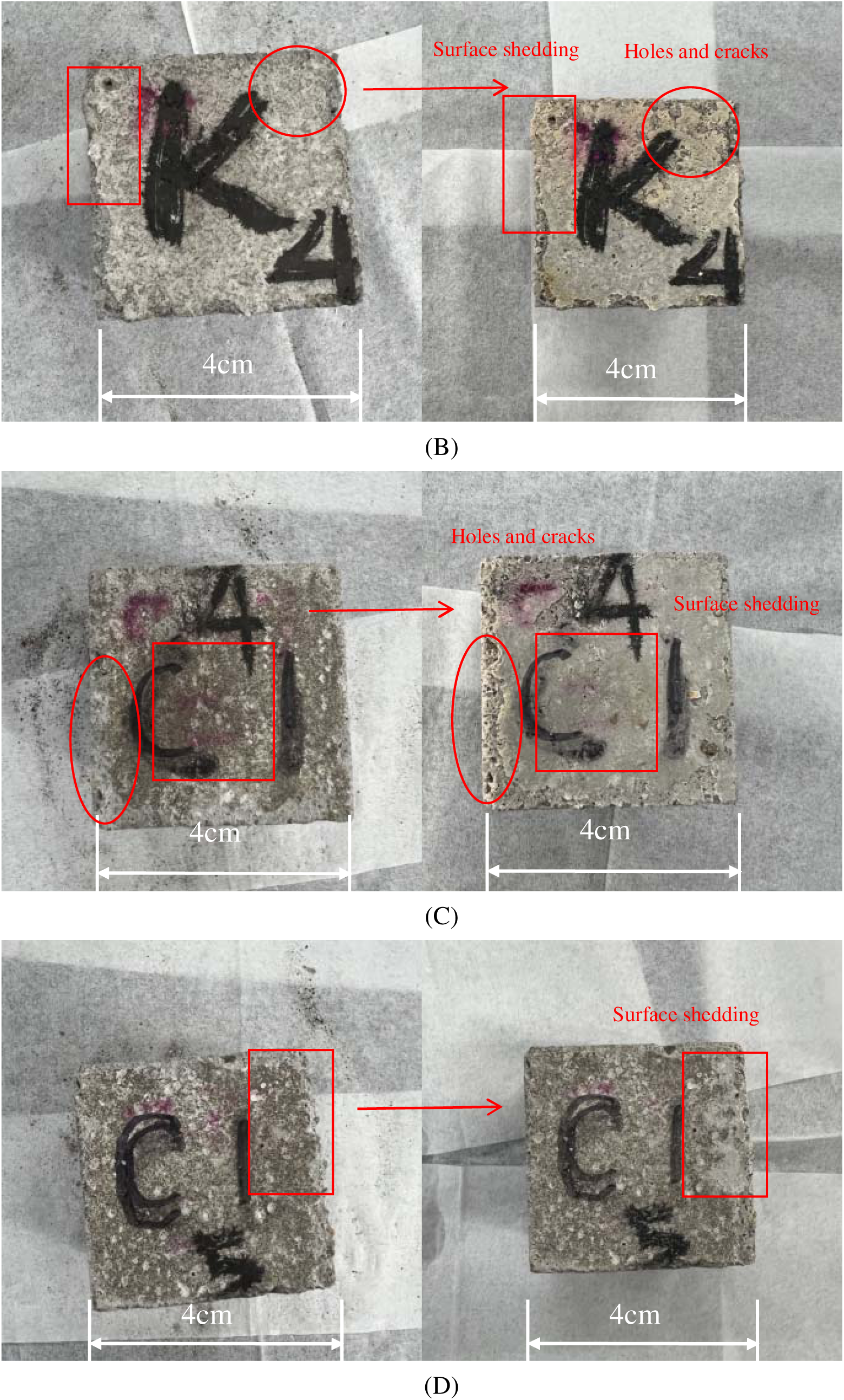
Top surface morphology of coupons before and after exposure. (A) Group K coupon before (left) and after 7 days of exposure (right), (B) Group K coupon before (left) and after 42 days of exposure (right), (C) C1 coupon before (left) and after 42 days of exposure (right), and (D) C1 coupon before (left) and after 56 days of exposure (right).

After 28 days of exposure, a yellow-colored compound was observed on the coupon side surface near the water level (Fig. S2). This observation could be attributed to the oxidation of H_2_S, leading to the formation of elemental sulfur. After 42 days of exposure, the surface of Group K coupons had developed a light-yellow discoloration and exhibited an increased density of cracks and pores (red circle box in Fig. 1B). Spalling of cenment mortar from the surface was observed, particularly near the edges of the coupon (red rectangle box in Fig. 1B).

In comparison, the coupon containing 0.05wt% nickel (N1) had little color change and less mortar spalling (Fig. S2). Meanwhile, coupons containing 0.05 wt% copper oxide (C1) had a lighter yellow surface stain compared to Group K, but still showed more cracks and pores (red circle box in Fig. 1C) and edges spalling (red rectangle box in Fig. 1C) than before exposure. After 56 days of exposure, spalling of cement mortar was clearly observed on the surface of C1 (Fig. 1D).

### 3.2 Top Surface pH

Prior to exposure, the top surface pH of the coupons was 10.06 ± 0.16, which was lower than that of freshly prepared cement mortar. This difference could be attributed to the required transportation and storage period of about 10 days. During this period, carbonization by atmospheric CO_2_ resulted in a pH decreas. After 7 days of exposure, the surface pH of all coupons increased to varying degrees, likely due the precipitation of alkaline substances from the coupon interior facilitated by accumulated surface water (Fig. 2). After 28 days of exposure, the top surface pH of Group K decreased sharply from 10.79 ± 0.06 to 7.51 ± 0.10 (Fig. 2A). The carbonization by CO_2_ typically reduces the surface pH of cementitious materials to about 10, but not to around 7, so the pronounced pH decrease in Group K was more likely attributed to the influence of an acidic substance, such as H_2_SO_4_, formed by the oxidation of H_2_S. During the subsequent exposure period from 28 to 56 days, the surface pH of Group K increased from 7.51 ± 0.10 to 10.37 ± 0.05. This rise may be explained by the ongoing acid corrosion, which continuously exposed the fresh alkaline material from the interior. During this process, the rate of alkaline material precipitation likely exceeded the corrosion rate, disrupting the dynamic equilibrium and leading to a sustained pH increase.

**Fig. 2.**
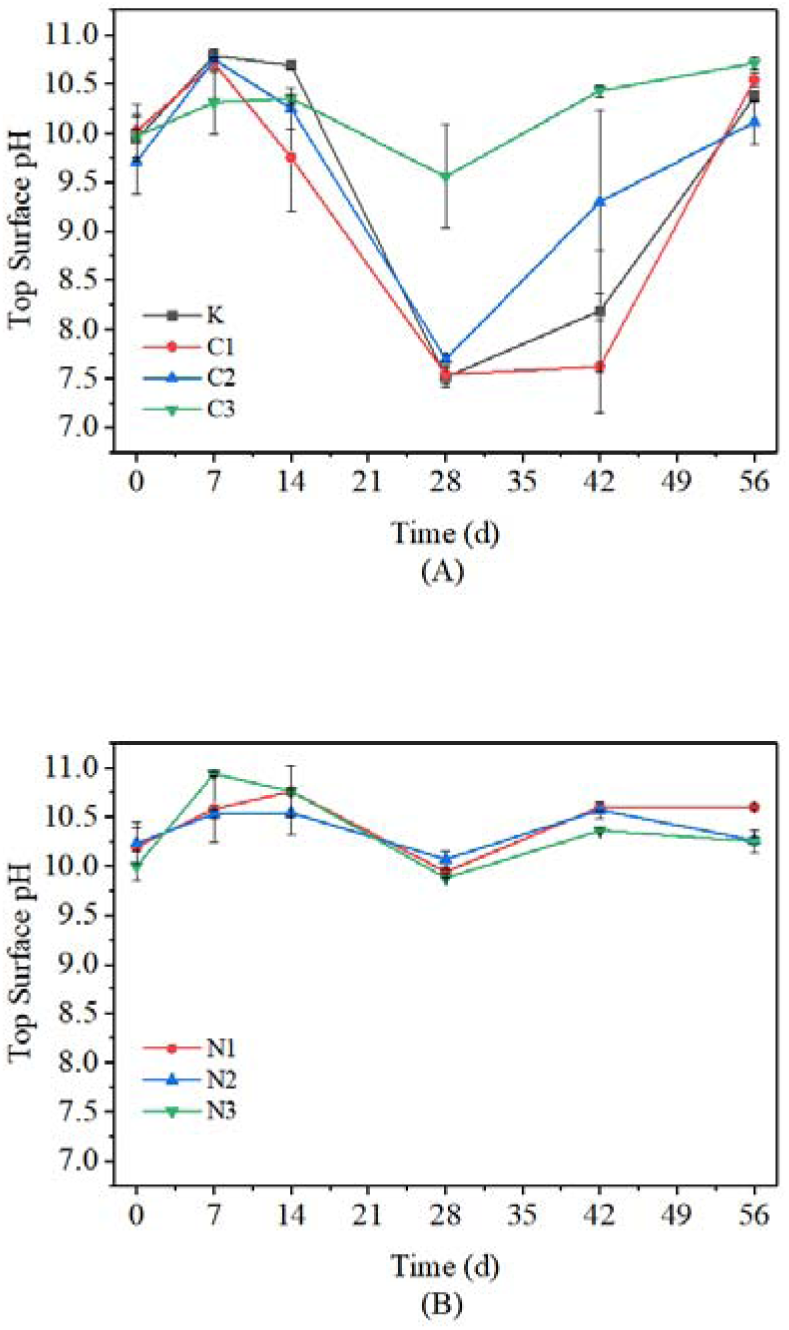
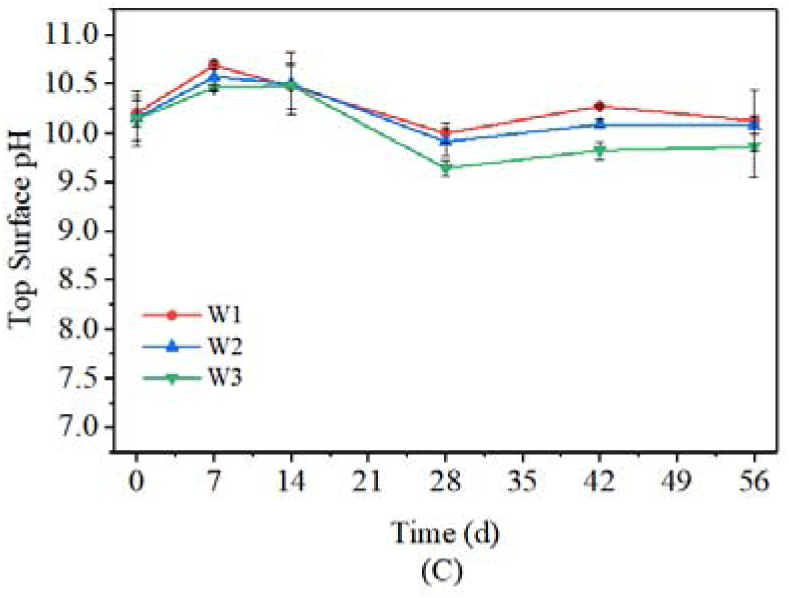
The top surface pH of (A) Group C, (B) Group N, and (C) Group W.

The trend in top surface pH variation observed in Group C, specifically for C1 and C2, closely resembled that of Group K. After 7 days of exposure, C1 and C2 exhibited increases in top surface pH to 10.71 ± 0.06 and 10.75 ± 0.01, respectively. Subsequently, after 28 days of exposure, their pH decreased to 7.54 ± 0.09 and 7.70 ± 0.04, respectively, before resuming an upward trend to reach 10.54 ± 0.07 and 10.11 ± 0.22 after 56 days of exposure. The pH variation of C3 exhibited a distinct trend. After 28 days of exposure, the pH of C3 remained at 9.56 ± 0.53, notably higher than that of C1, C2, and Group K. This trend persisted throughout the 28 to 56 days exposure period, with C3 consistently maintaining a higher pH than the other groups.

The overall trend in top surface pH change observed in both Group N and Group W closely resembled that of Group C3 (Fig. 2B). After 7 and 14 days of exposure, the pH of N1 increased to 10.58 ± 0.33 and 10.76 ± 0.04, respectively. After 28 days, it decreased to 9.94 ± 0.02 which was higher than that of Group C and Group K, then increased to 10.60 ± 0.02 after 56 days which remained higher than Group C and K. The pH change trend for N2 and N3 closely followed that of N1, with all these coupons maintaining higher pH than Group C and K during the 28- and 42-day exposure period. The primary difference emerged after 56 days of exposure, when the pH of N2 and N3 measured 10.26 ± 0.04 and 10.25 ± 0.11, respectively, which was slightly lower than that of N1.

The top surface pH trend observed in Group W was similar to that in Group N, with both groups maintaining higher pH than C1, C2 and K during the 14 to 56 days of exposure (Fig. 2C). However, after 28 days of exposure, the pH of W1was 10.00 ± 0.10, exceeding that of W2 and W3 (9.91 ± 0.14 and 9.64 ± 0.08, respectively). This difference persisted after 56 days of exposure, with pH of W1 measuring 10.12 ± 0.31, again higher than W2 and W3 (10.08 ± 0.09 and 9.86 ± 0.31, respectively). Throughout the period from 28 to 56 days of exposure, the pH of W1consistently exceeded that of W2 and W3.

### 3.3 Side surface pH

After 28 days of exposure, the side surface pH of Group K decreased from 10.33 ± 0.06 to 9.99 ± 0.22 (Fig. 3A). This decrease might be attributed to the lower pH of the sewage (7.21 ± 0.13) compared to the initial pH of the cement mortar. The submerged portion of the coupons was directly affected by the sewage, resulting in a slight pH reduction. After 56 days of exposure, the pH of Group K exhibited a further decrease to 10.13 ± 0.60. This fluctuation may reflect the dynamic balance between acid corrosion (or sewage immersion) after submerged part of the coupon and the precipitation of alkaline substances within the coupon matrix.

**Fig. 3.**
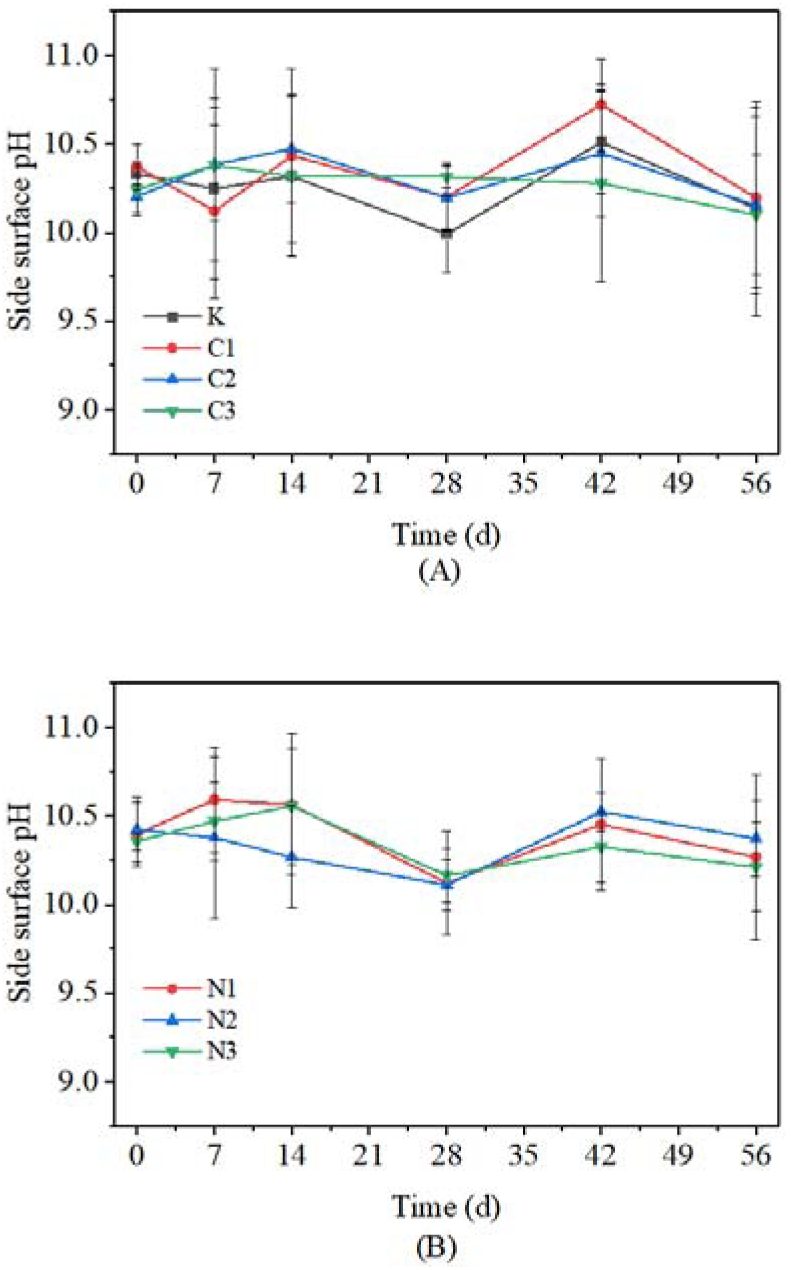
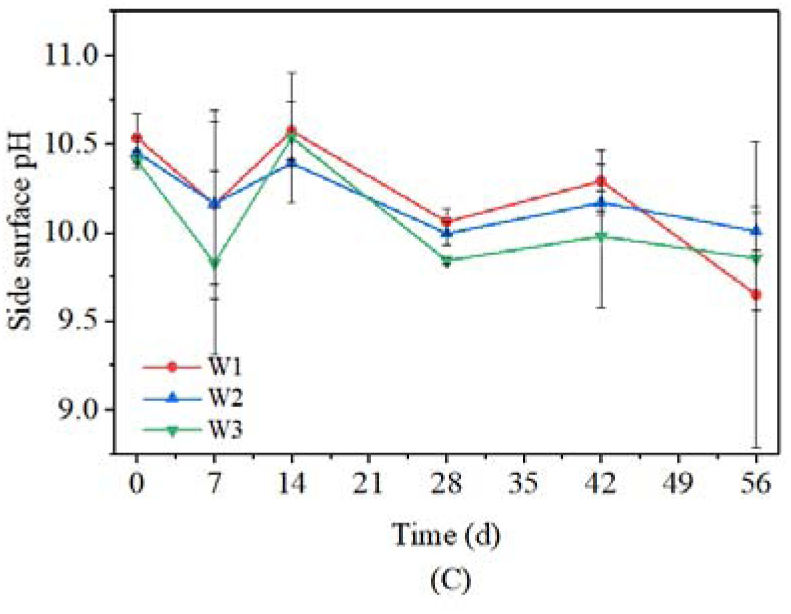
Side surface pH of different coupon groups. (A) Group C, (B) Group N, and (C) Group W.

Compared to Group K, the side surface pH of Group C exhibited a similar fluctuation trend over the 56-day exposure period (Fig. 3A). The pH fluctuations of C2 and C3 were not significantly different from those of Group K during this time frame. However, a notable distinction emerged after 42 days of exposure, when the side surface pH of C1 increased to 10.72 ± 0.26, surpassing that of K (10.51 ± 0.29), C2 (10.45 ± 0.36), and C3 (10.28 ± 0.56) during the same period. Notably, the pH of C1 consistently measured higher than that of C2, C3, and K between days 14 and 56 of exposure.

Group N also displayed a pattern similar to that of Group K: after 28 days of exposure, the pH of N1, N2, and N3 decreased to 10.12 ± 0.30, 10.11 ± 0.14, and 10.16 ± 0.15, respectively, and subsequently rose to 10.27 ± 0.47, 10.37 ± 0.21, and 10.21 ± 0.25, respectively, after 56 days of exposure (Fig. 3B). In the case of Group W (W1/W2/W3), the surface pH decreased to 10.06 ± 0.07, 9.99 ± 0.07, and 9.84 ± 0.02, respectively, after 28 days of exposure (Fig. 3C). After 56 days of exposure, the pH further decreased to 9.65 ± 0.86, 10.01 ± 0.11, and 9.85 ± 0.29, falling below the levels observed in Group K.

### 3.4 Sulfate concentration

The changes of sulfate concentration in the corrosion layer of coupons are depicted in Figure 4. The sulfate concentration of Group K increased over time, from 0.0025 mg/cm^2^ to 10.65 mg/cm^2^ after 56 days of exposure (Fig. 4A).

**Fig. 4.**
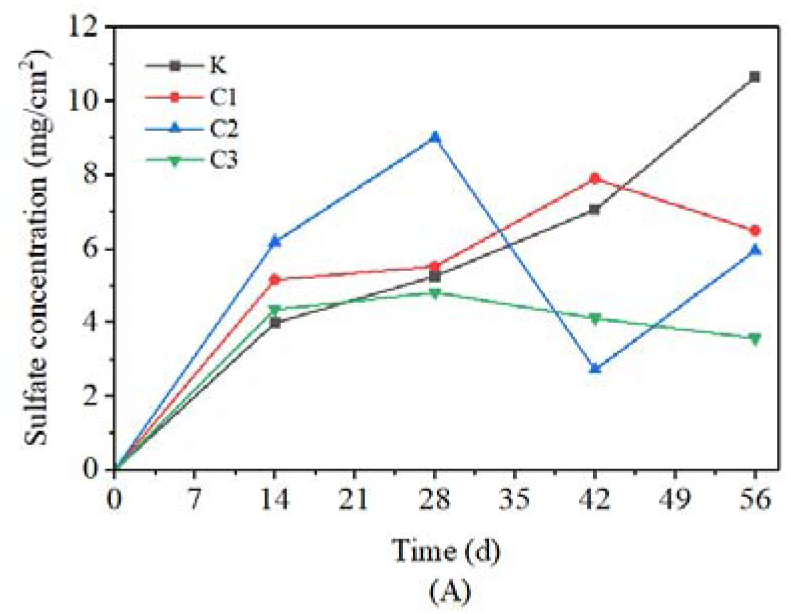
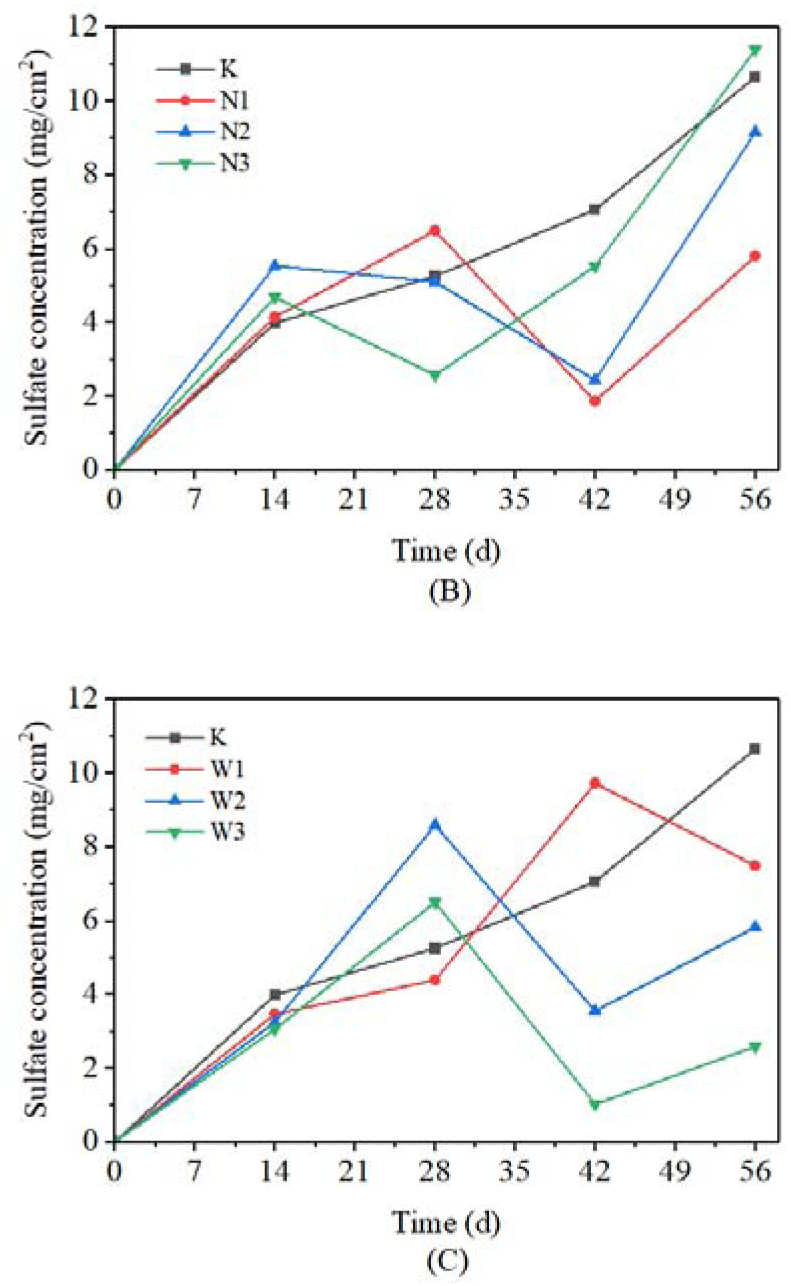
The sulfate concentration of (A) Group C, (B) Group N, (C) Group W in corrosion layer during 56 days of exposure.

During the 28-day exposure period, the sulfate concentration of Group C followed a similar trend to that of Group K, increasing to 5.51 mg/cm^2^, 9.01 mg/cm^2^, and 4.81 mg/cm^2^, respectively. However, in the subsequent period, C2 and C3 exhibited a decline. After 42 days of exposure, they decreased to 2.70 mg/cm^2^ and 4.1 0mg/cm^2^, respectively, while C1 continued to increase to 7.05 mg/cm^2^. After 56 days of exposure, C1 and C3 decreased to 6.47 mg/cm^2^ and 3.56 mg/cm^2^, respectively, while C2 increased to 5.94 mg/cm^2^ (Fig. 4A).

After 14 days of exposure, the sulfate concentration of N1, N2 and N3 increased to 4.14 mg/cm^2^, 5.53 mg/cm^2^ and 4.67 mg/cm^2^, respectively (Fig. 4B). After 28 days of exposure, the sulfate concentration of N1 increased to 6.48 mg/cm^2^, and those of N2 and N3 decreased to 5.09 mg/cm^2^ and 2.56 mg/cm^2^, respectively. After 42 days of exposure, the sulfate concentration of N1 and N2 decreased to 1.86 mg/cm^2^ and 2.42 mg/cm^2^, respectively. However, the sulfate concentration of N3 exhibited a continuous increase, reaching 5.51 mg/cm^2^ and eventually 11.39 mg/cm^2^ after 56 days of exposure.which is slightly higher than that of K (10.65 mg/cm^2^). And the sulfate concentration of N1 and N2 increased to 5.79 mg/cm^2^ and 9.15 mg/cm^2^, respectively, after 56 days of exposure, which were lower than that of K.

The changes in sulfate concentration of W2 and W3 showed a similar trend to C2, rising continuously to 8.59 mg/cm^2^ and 6.49 mg/cm^2^ after 28 days of exposure, then declining to 3.54 mg/cm^2^ and 1.02 mg/cm^2^, respectively, after 42 days of exposure, and finally rising to 5.81 mg/cm^2^ and 2.58 mg/cm^2^, after 56 days of exposure (Fig. 4C). The change trend of W1 was similar to C1, first rising to 9.71 mg/cm^2^ after 42 days of exposure and then decreasing to 7.48 mg/cm^2^ after 56 days of exposure.

### 3.5 Mass changes

The mass of all coupons after exposure in the corrosion chamber was higher than those prior to exposure (Fig. 5). R of Group K showed a trend of first increase and then decrease. After 14 days of exposure, R of Group K increased from 0% to 1.35%, and then the growth slowed down, increased to 1.54% after 28 days of exposure. The increase in R may be attributed to continuous sewage absorption when the coupons were immersed in sewage, leading to increased water content within the coupons. Additionally, the coupons were situated in a high-humidity environment, resulting in the accumulation of condensation water on the surface. It was also possible that chemical or biological oxidation has converted H_2_S into SO_4_^2-^ or S, which has subsequently accumulated on coupons resulted to mass increase. After 42 days of exposure, R decreased slightly to 1.49%. After 56 days, it dropped to 0.49%. After 42 days of exposure, partial shedding of the coupon surface occurred (Figure 1C and 1D). Hence, the decline of R may be attributed to an imbalance between the accumulation of materials on the coupons and the surface shedding of the coupons surface during the exposure period.

**Fig. 5.**
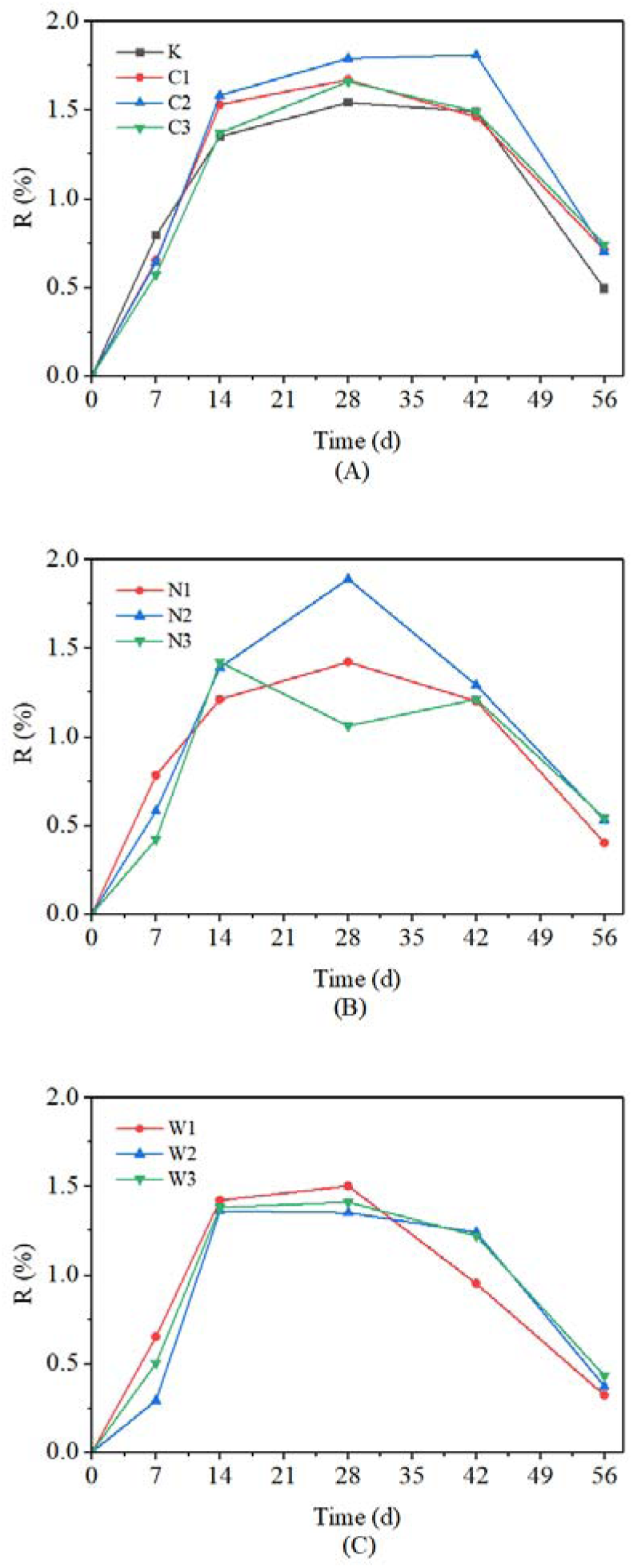
The R of (A) Group C, (B) Group N, (C) Group W during 56 days of exposure.

After 28 days of exposure, the R of Group C (C1, C2, and C3) increaced to 1.67%, 1.79%, and 1.66%, respectively, higher than K (1.54%). Subsequently, the change trend of R of C1 and C3 mirrored that of K, exhibiting a decreasing trend, and after 56 days of exposure, were decrease to 0.71% and 0.74%, respectively, slightly higher than K (0.49%). However, after 42 days of exposure, R for C2 continued to increase to 1.81%, and after 56 days of exposure, it subsequently decreased to 0.7%.

The R of N1 and N3 after 28 days of exposure were 1.42% and 1.06%, respectively, which were lower than K (1.54%). However, the R of N2 was 1.89% whitch higher than K after 28 days of exposure. During 42-56 days of exposure, the R of N1, N2, and N3 were lower than K and Group C.

During the exposure period, the trend of R change in W2 and W3 closely resembled that of K. W1 exhibited a higher rate of R increase than W2 and W3 within the 28 days of exposure. However, after 42 days of exposure, the R of W1 rapidly decreased to 0.95%, lower than W2 and W3 (1.24% and 1.22%, respectively). After 56 days of exposure, the R of W1 decreased to 0.32%, which was slightly lower than W2 and W3 (0.37% and 0.43%, respectively).

### 3.6 Characterization of the corrosion layer by Scanning Electron Microscopy and Energy Dispersive X-ray Spectroscopy

The SEM analysis of the corrosion layer, exposed to the gas-phase environment, was performed on coupons both before exposure (Group K) and after 56 days of exposure (Group K, C3, N1, N3, and W3)

Microcracks in different sizes were observed in all SEM images.SEM images of Group K before exposure were depicted in Figure 6A. Dark gray irregular substances (indicated by red circle 1 in Fig. 6A) were observed, surrounded by light grayish-white substances (indicated by red circle 2 in Fig. 6A). The EDS analysis revealed that the dark gray irregularly shaped region had the highest Si and O content (see Figure S3A2). This suggested the presence of SiO_2_, which are consistent with the standard sand used in the preparation of cement mortar. Additionally, the substance surrounding the SiO_2_ exhibited the highest content of Ca, along with traces of C, Mg, Al, and Si, indicating that it may be the hydration product of cement, specifically, calcium silicate hydrate (C-S-H).

**Fig. 6.**
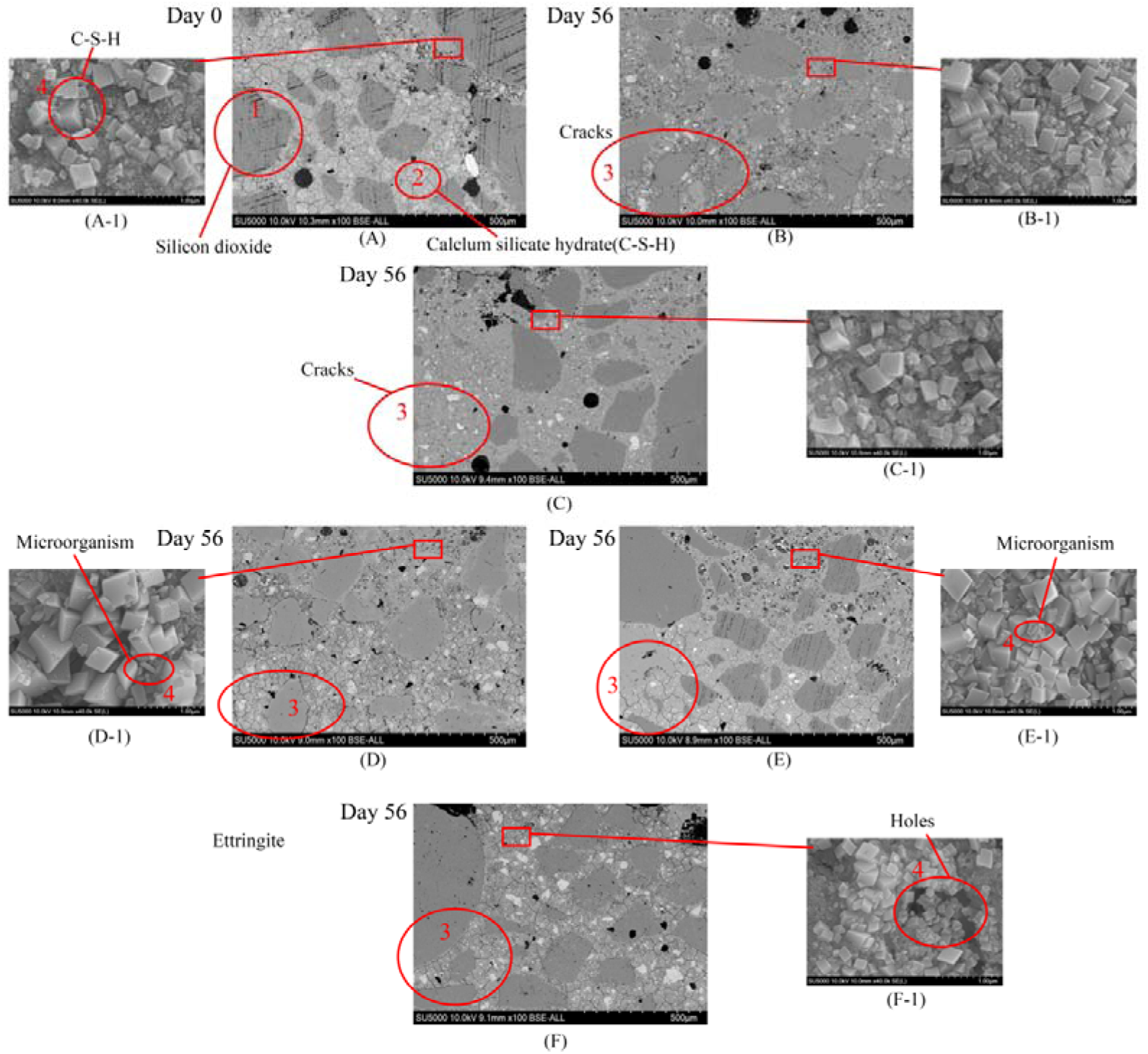
SEM images of corrosion layer at the top surface of Group K (A) before and (B) after 56 days of exposure, respectively. (C), (D), (E), and (F) are SEM images of C3, N1, N3, and W3 after 56 days of exposure, respectively. A-1, B-1, C-1, D-1, E-1, and F-1 are magnified SEM images of the regions pointed by red line and empty red circle.

Fig. 6B presented the SEM image of Group K after 56 days of exposure. In comparison to the pre-exposure state, the post-exposure image reveals the presence of more micro-cracks (highlighted by red circle 3 in Figure 6B). Further analysis through EDS demonstrates that, after 56 days of exposure, aluminum (Al) in the K corrosion layer becomes increasingly concentrated, while calcium (Ca) disperses from the concentrated region to the surrounding area (refer to Figure S3B2). Table S3 indicates an overall increase in the total sulfur (S) content in the sample. These elemental changes suggest that after 56 days of exposure, calcium within the C-S-H dissociates and combines with aluminum and sulfur to form concrete corrosion products such as ettringite (3CaO•Al_2_O_3_•3CaSO_4_•32H_2_O) and gypsum (CaSO_4_• 2H_2_O).

Figure 6C showed SEM images of C3 after 56 days of exposure. These images revealed significant cracks developed on the coupon surface (red circle 3 in Figure 6C). Additionally, the sulfur (S) distribution on the surface of C3 appears more dispersed, which could indicate an increase in sulfur content within the coupon. However, no significant sulfur enrichment was observed in regions of calcium (Ca) and aluminum (Al) enrichment. This observation suggested that Ca, Al, and S were not strongly bound within the coupon and there may not be a significant amount of corrosion products, such as gypsum and ettringite, formed on the coupon (Figure S3B-1). Similar to the C3, N1, N3, and W3 also exhibited varying degrees of cracking after 56 days of exposure, as depicted by red circles 3 in Figures 6D, 6E, and 6F. Upon examining the EDS images, it became apparent that aluminum (Al) and sulfur (S) in N1 were more dispersed compared to N3. This dispersion could suggest that calcium (Ca) has detached from C-S-H and combined with Al and S to form corrosion products like gypsum and ettringite (Figures S3D and E). Conversely, the W3 contained higher sulfur and aluminum content, and sulfur clusters alongside aluminum and calcium. This clustering may indicate that the W3 sample has accumulated more corrosion products, relative to other coupons (Figure S3F).

To further understand the microscopic changes in coupon surface, the area of interest (red rectangular boxed area in Figure 6) was analyzed with higher magnification SEM. Upon magnified view (Figures A-1, B-1, C-1, D-1, E-1), it was observed that almost all coupons contained distinct rectangular or quadrilateral crystals (red circle 4 in Figure 6A-1). The presence of this crystal in Group K both before and after exposure, combined with the material composition of the enlarged area, suggests that this crystal may indeed be C-S-H. On the surfaces of N1 and N3, as well as within the crevices, small cylindrical or rod-like structures were detected, which are likely to be microorganisms (as indicated by the red circle 4 in Figure 6D-1 and E-1). Concurrently, there was a noticeable widening of gaps within the C-S-H structure in W3, leading to the gradual formation of holes (as depicted by the red circle 4 in Figure 6F-1).

### 3.7 ATP content

The ATP content of K increased from 0 to 80.21*10^−6^ ± 3.51*10^−6^ μmol/cm^2^ after 14 days of exposure (Fig. 7). Subsequently, it exhibited a fluctuating trend and decreased to 36.81*10^−6^ ± 3.19 *10^−6^ μmol/cm^2^ after 28 days of exposure. After 42 days of exposure, it increased to 54.98*10^−6^ ± 6.18 *10^−6^ μmol/cm^2^ followed by a decrease to 30.17*10^−6^ ± 3.87 *10^−6^ μmol/cm^2^ after 56 days of exposure.

**Fig. 7.**
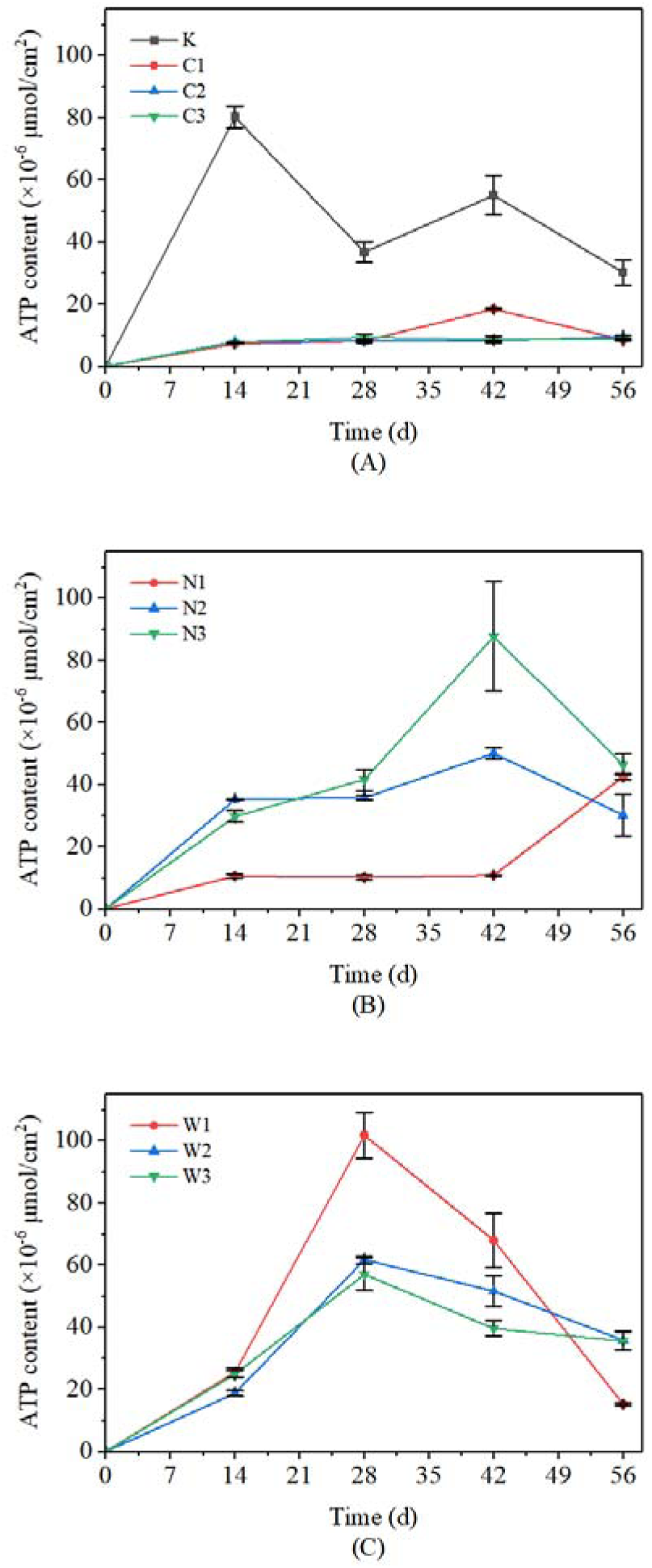
The ATP content changes of (A) Group C, (B) Group N, (C) Group W in the corrosion layer during 56 days of exposure.

During the exposure period, the ATP content of all coupons in Group C was significantly lower than that K. After 14 days of exposure, the ATP content of C1, C2, and C3 increased from 0 to 7.19*10^−6^ ± 0.13 *10^−6^ μmol/cm^2^, 7.88 ± 0.01 *10^−6^ μmol/cm , and 7.71*10 ± 0.17 *10 μmol/cm , respectively, which was much lower than that of K (80.21*10^−6^ ± 3.51 *10^−6^ μmol/cm^2^). The ATP content of C2 and C3 remained relatively stable from 14 to 56 days of exposure, measuring 9.44*10^−6^ ± 0.63 *10^−6^ μmol/cm^2^ and 8.64*10^−6^ ± 0.38 *10^−6^ μmol/cm^2^, respectively, after 56 days of exposure. However, the ATP content of C1 increased to 18.42*10^−6^ ± 0.28 *10^−6^ μmol/cm^2^ after 42 days of exposure and then decreased to 8.51*10^−6^ ± 0.50 *10 μmol/cm^2^ after 56 days of exposure.

The changeing trend of ATP content in N1 was similar to Group C. After 14 days of exposure, it increased from 0 to 10.58*10^−6^ ± 0.58 *10^−6^ μmol/cm^2^, and then remained relatively stable until it reached 10.77*10^−6^ ± 0.13 *10^−6^ μmol/cm^2^ after 42 days of exposure. It further increased to 42.46*10^−6^ ± 0.96 *10^−6^ μmol/cm^2^ after 56 days of exposure, with the ATP content of N1 consistently lower than K during the exposure period. After 14 days of exposure, the ATP content of N2 and N3 increased to 35.1*10^−6^5 ± 0.18 μmol/cm^2^ and 29*10^−6^.73 ± 1.91 *10^−6^ μmol/cm^2^, respectively, and then to 49.98 ± 1.86 *10^−6^ μmol/cm^2^ and 87.67 ± 17.54 *10^−6^ μmol/cm^2^, respectively, after 42 days of exposure. During this period, the ATP content of the N2 and N3 exceeded Group C and N1. However, after 56 days of exposure, the ATP content of N2 and N3 decreased to 30.17*10^−6^ ± 6.74 *10^−6^ μmol/cm^2^ and 46.38*10^−6^ ± 3.58 *10^−6^ μmol/cm^2^, respectively.

During the exposure period, the ATP content of group W exhibited an initial increase followed by a decrease. After 28 days of exposure, the ATP content of W1, W2, and W3 increased to 101.71*10^−6^ ± 7.45 *10^−6^ μmol/cm^2^, 61.67*10^−6^ ± 1.15 *10^−6^ μmol/cm^2^ , and 56.94 ± 5.19 *10 μmol/cm , respectively, and then decreased to 15.25*10^−6^ ± 0.44 *10^−6^ μmol/cm^2^, 35.72*10^−6^ ± 3.16 *10^−6^ μmol/cm^2^, and 35.5*10^−6^ ± 2.84 *10^−6^ μmol/cm^2^, respectively, after 56 days of exposure.

## 4. Discussion

### 4.1 The corrosion process of coupons

Throughout the exposure process, the surface appearance, surface pH, mass, sulfate concentration, and ATP content exhibited dynamic changes with varying exposure times. After 14 days of exposure, the side surface pH of Group K decreased from 10.33 ± 0.06 to 10.32 ± 0.45 (Fig. 3A), while the top surface pH increased from 9.94 ± 0.22 to 10.69 ± 0.04 (Fig. 2A). The decrease in pH on the side surface can be attributed to side surface in direct contact with the sewage. The pH of the sewage used was 7.16 ± 0.08, which was lower than the surface pH of coupons, leading to a mild acidification effect on the side surface pH. Notably, after 14 days of exposure to high humidity (100% RH), the surface of Group K became wetter compared to their pre-exposure state. Given the uneven surface of coupons, water accumulated significantly in the depressions of the top surface (Fig. 1A). This accumulation of water on the top surface triggered the precipitation of alkaline substances within the cement mortar, resulting in an increase in the top surface pH. Furthermore, during this period, the ATP content increased to 80.21*10^−6^ ± 3.51*10^−6^ μmol/cm^2^ (Fig. 7A). ATP content was non-selective and do not directly indicate the total amount of acid-producing microorganisms within the coupons. At this stage, the surface pH of coupons still maintained a slightly alkaline environment (pH > 10), which was less conducive to the colonization of acid-producing microorganisms [10]. Therefore, the rapid rise in ATP content may be attributed to the coupon’s immersion in sewage, allowing microorganisms present in the sewage to adhere to the coupon surface. After 14 days of exposure, the sulfate concentration increased from 0.0025 mg/cm^2^ to 3.98 mg/cm^2^ (Fig. 4A), indicating sustained sulfate input to Group K. The accumulation of sulfate on the coupon surface can be attributed to both chemical oxidation and microbial oxidation processes [41]. However, the surface pH of coupon was not conducive to colonization by acid-producing microorganisms at this stage, as mentioned earlier. Research has suggested that the chemical oxidation rate of H_2_S is positively correlated with humidity [42]. Therefore, the most likely cause of the increase in sulfate content is the chemical oxidation process of H_2_S. This accumulation of sulfate also resulted in an increase in the sample’s mass (Fig. 5).

After 28 days of exposure, the top surface pH of Group K decreased from 10.69 ± 0.04 to 7.51 ± 0.10 (Fig. 2A), while the side surface pH only decreased from 10.32 ± 0.45 to 9.99 ± 0.22 (Fig. 3A). The alteration in surface pH could be attributed to two primary reasons: (1) The pH reduction due to the influence of acidic substances; (2) Increased surface water content, leading to the precipitation of alkaline substances within the cement mortar and resulting in an increase in pH. Between days 14 and 28 of exposure, the top surface pH of Group K experienced a significant drop, which may be attributed to coupon corrosion caused by the presence of acidic substances. As the surface pH dipped below 9, the colonization of neutrophilic sulfur oxidizing bacteria (NSOB) commenced on the coupon’s surface [43]. This colonization led to an intensification of microbial oxidation of H_2_S, resulting in increased biological acid production and a rapid decrease in surface pH. Concurrently, the rate of precipitation of alkaline substances within the sample was influenced. During this period, the sulfate concentration of Group K increased from 3.98 mg/cm^2^to 5.25 mg/cm^2^ (Fig. 4A). However, the ATP content decreased form 80.21*10^−6^ ± 3.51*10^−6^ μmol/cm^2^ to 36.81*10^−6^ ± 3.19 *10^−6^ μmol/cm^2^ (Fig. 7A). This decline in ATP content could be attributed to the initial stage of corrosion, where microbial species on the coupon surface were still changing, and the total population of acid-producing microorganisms was relatively low. It’s noteworthy that, during this phase, there were observations of the accumulation of a small quantity of yellow material on the coupon surface and partial surface shedding (Figure S2A). This yellow substance is reported to potentially be elemental sulfur (S) resulting from the oxidation of H_2_S [44]. The combined accumulation of microorganisms and H_2_S oxidation products led to an increase in R. However, the surface layer shedding resulted in a loss of mass for the coupon, ultimately slowing down the rate of R growth.

After 42 days of exposure, the top surface pH of Group K exhibited a rapid increase. This upward trend persisted until 56 days of exposure, ultimately reaching 10.37 ± 0.05 (Fig. 2A). Simultaneously, the sulfate concentration continued to increase from to 7.05 mg/cm^2^ (after 42 days of exposure) and 10.65 mg/cm^2^ (after 56 days of exposure) , while the ATP content initially rose to 54.98*10^−6^ ± 6.18*10^−6^ μmol/cm^2^ (after 42 days of exposure) and then declined to 30.17*10 ± 3.87*10 μmol/cm (after 56 days of exposure). Throughout the exposure period from 28 to 56 days, both the sulfate concentration and ATP content displayed fluctuations. Following the classical three-stage corrosion model, when the concrete surface pH falls within the range of 9 to 6, various acid-producing microorganisms, such as *Thiobacillus thiooxidans, Starkeya novella, Halothiobacillus neopolitanus*, and *Thiobacillus intermedius*, colonize the concrete surface [43]. Subsequently, H_2_S is further oxidized into sulfuric acid, leading to concrete corrosion . Therefore, during the exposure period from 28 to 56 days, the types and quantities of NSOB continued to increase, resulting in the production of more biological acids as the surface pH of the coupons steadily declined. This period also saw the emergence of noticeable holes and cracks on the surface of Group K, along with the shedding of the surface layer (Fig. 1B). SEM and EDS results confirmed the presence of corrosion products like ettringite (3CaO• Al_2_O_3_ • 3CaSO_4_ • 32H_2_O) and gypsum (CaSO_4_• 2H_2_O) (Figure 6, Figure S3). As the corrosion process advanced, the overall population of microorganisms on the coupon surface increased. However, surface shedding also removed some microorganisms and H_2_S oxidation products. Consequently, the rate of mass accumulation on the coupon decreased, leading to a gradual decline in R (Fig. 5A).

The side surface pH of the coupons during the exposure period exhibited relatively minor fluctuations compared to the top surface pH, displaying an overall trend of wavering changes. This difference might be attributed to the immersion of the coupons in sewage, where the side surface pH was influenced by the sewage pH. Throughout the exposure duration, the sewage was replaced every 14 days, and over this period, the sewage pH increased from 7.16 ± 0.08 to 9.58 ± 0.32. This suggested that the coupons immersed in sewage continued to release alkaline substances into the sewage throughout the exposure.

In summary, the corrosion progression of Group K can be roughly delineated into three stages:

Stage 1 (0-14 days of exposure): During this initial stage, the coupons were influenced by environmental factors such as CO_2_, H_2_S, and humidity [42, 44]. The surface pH of the coupons decreased, and the chemical oxidation of H_2_S led to the accumulation of sulfate concentration. Additionally, microbial activity in the sewage contributed to an increase in ATP.

Stage 2 (14-28 days of exposure): In this stage, the surface pH of coupons dropped to levels conducive to colonization by acid-producing microorganisms. This condition promoted the conversion of sulfur elements into sulfate through microbial and chemical oxidation processes. The corrosion continued to progress, resulting in the formation of corrosion products like ettringite (3CaO•Al_2_O_3_•3CaSO_4_•32H_2_O) and gypsum (CaSO_4_•2H_2_O). As a certain amount of corrosion products accumulated, the coupon’s surface layer began to shed. This shedding process removed some surface-layer microorganisms and H_2_S oxidation products (SO_4_^2-^ and S), leading to a decline in both R and ATP content.

Stage 3 (28-56 days of exposure): During the corrosion process in this stage, the development of corrosion within the corrosion layer and the ongoing surface layer shedding became dynamic processes. This led to fluctuations in surface pH, sulfate concentration, and ATP content.

### 4.2 Inhibition effect of different metal bacteriostatic agent additives on corrosion process

The corrosion development process in Group C, Group N, and Group W during exposure followed a similar pattern to Group K and could be roughly categorized into three stages.

Stage 1: After 14 days of exposure, there was a variable increase in both the top surface pH and R of all coupons, with minimal variation between the different groups. At this point, the average surface pH of Group C, Group N, and Group W were 10.12 ± 0.32, 10.69 ± 0.13, and 10.48 ± 0.02, respectively, which did not significantly differ from those of Group K (10.69 ± 0.04). However, it’s important to note that during this stage, the surface pH of all samples remained above 9, which is not conducive to the colonization of acid-producing microorganisms. Consequently, corrosion development during this period was primarily driven by the chemical oxidation of H_2_S. Therefore, the use of the metal bacteriostatic agent did not appear to have a significant impact on corrosion development during this stage.

Stage 2: After 28 days of exposure, the top surface pH of C1 and C2 experienced a rapid decrease to 7.54 ± 0.09 and 7.70 ± 0.04, respectively, exhibiting a similar declining trend as Group K (decrease from 10.69 ± 0.04 to 7.51 ± 0.10). This change made them gradually suitable for colonization by NSOB. However, the top surface pH of C3 only decreased from 10.35 ± 0.04 to 9.56 ± 0.53, maintaining relatively alkaline conditions. During this period, the sulfate concentration of C3 increased to 4.81 mg/cm^2^, which was lower than that of C1 (5.51 mg/cm^2^), C2 (9.01 mg/cm^2^) and K (5.25 mg/cm^2^). This indicates that the H_2_S oxidation process on the surface of C3 was less intense compared to C1, C2 and Group K. Since all coupons were immersed in sewage with a relatively constant sulfate concentration (61.38±0.93 mg/L), the increase in sulfate on the coupon surface over time appeared to be approximately linear. Therefore, the rise in sulfate concentration on the coupon surface was likely due to chemical and/or microbial oxidation of H_2_S, rather than being influenced by sulfate present in the sewage. As all coupons were exposed under the same environmental conditions, their chemical oxidation rates were not different. Consequently, the variation in sulfate concentration on coupon surfaces was more likely driven by microbial oxidation. Additionally, the ATP content and R of C3 were lower than those of C2, C3 and Group K, indicating a lower total number of microorganisms on the surface of C3. The sulfate concentration and ATP content results together demonstrated that a higher amount of copper oxide added had a better inhibitory effect on H_2_S microbial oxidation during the second stage of corrosion.

After 28 days of exposure, the ATP content of N1 (10.12×10^−6^ ± 0.78×10^−6^ μmol/cm^2^ ) was lower than that of N2 (35.81×10^-6^ ± 0.88×10^-6^ μmol/cm ) and N3 (29.73×10^−6^ ± 1.91×10^−6^ μmol/cm^2^). However, the sulfate concentration of N1 was higher than N2 and N3. This difference may be attributed to the inhibitory effect of lower nickel content on H_2_S microbial oxidation, resulting in a delayed corrosion process for N1. Moreover, N2 and N3 exhibited higher ATP content, indicating more active acid-producing microorganisms on the surfaces, leading to increased production of biological acids during corrosion. This, in turn, led to the formation of corrosion products like gypsum and ettringite. However, in high humidity environments, H_2_S oxidation products like sulfate are more likely to be lost along with the surface layer of the coupon. When the surface layer sheds, R also decreases (Fig. 5B). Consequently, the sulfate concentration of N2 and N3 were lower than N1. Additionally, the surface shedding observed in N2 and N3, along with the presence of more gaps and holes on the surface, supports this hypothesis (Fig. S2 E, F, G)

Stage 3: After 28 days of exposure, the ATP content of W1 was higher than that of W2, W3, C3, N1 (Fig. 7). This suggested that the total number of microorganisms on the surface of W1 was relatively high. The intense microbial activity oxidizes more H_2_S to sulfate, resulting in a higher sulfate concentration in W1 compared to W2 and W3 (Fig. 4). As corrosion progresses, part of the surface layer of W1 sheds, leading to a rapid decrease in R of Group W (Fig. 5C). Simultaneously, the underlying alkaline cement mortar was exposed, causing an increase in surface pH (Fig. 2C). After 56 days of exposure, the top surface pH of W1 was higher than that of W2 and W3. During the 28 to 56 days of exposure, the W3 with the lowest ATP content and the lowest sulfate concentration in Group W surpassed the Group C and Group N with higher contents. This could be attributed to more microbial activity on the surface of Group W, leading to increased biological acid production. With the accumulation of corrosion products, the surface layer of Group W experienced more shedding, carrying away some of the accumulated corrosion products. This resulted in a greater decline in R of Group W compared to Group N and Group C (Fig. 5).

Therefore, three fundamental conclusions can be derived:

1. In the first stage of corrosion development, chemical oxidation of H_2_S primarily drives the process, and the inhibitory effects of different metal bacteriostatic agents on the biological oxidation of H_2_S do not exhibit significant differences at this stage.
2. In the second stage, afier 28 days of exposure, the surface pH, sulfate concentration, and ATP content of C3 (9.56 ± 0.53, 4.81 mg/cm^2^, and 9.31×10^−6^ ± 0.86×10^−6^ μmol/cm^2^, respectively,) closely resemble those of N1 (9.94 ± 0.02, 6.48 mg/cm^2^, and 10.12×10^−6^ ± 0.78×10^−6^ μmol/cm^2^, respectively,). This indicates that C3 and N1 exhibit superior inhibition of H_2_S biooxidation compared to other coupons of the same type with different dosages. The surface pH, sulfate concentration, and ATP content of W3 are 9.64 ± 0.08, 6.49 mg/cm^2^, and 56.94×10^−6^ ± 5.19×10^−6^ μmol/cm^2^, respectively, which are not superior to those of W1 and W2.
3. In the third stage, W3, with the lowest ATP content and the lowest sulfate concentration in Group W, surpasses that in Group C and Group N. This suggests, to some extent, that in the initial stage of corrosion, sodium tungstate exhibits a weaker inhibitory effect on H_2_S biological oxidation compared to copper oxide and nickel oxide.

Different types of metal bacteriostatic agents indeed exhibit distinct inhibition effects on MICC, which can be closely linked to pH and microbial species [32, 33, 36]. MICC is a long-term process, and the pH of sewer system concrete can decrease from 11 to approximately 1 over several years or even decades [45]. Throughout this process, the types and quantities of microorganisms on the concrete surface can undergo significant changes [43]. Nickel and tungsten can bind with *T.thiooxidans* NB1-3 and *A.thiooxidans*, respectively, to inhibit the activities of sulfur dioxygenase and sulfate oxidase, thereby inhibiting microbial growth [32, 34, 46, 47]. However, *T.thiooxidans* NB1-3 are suited for colonization in neutral environments (pH=7 ± 1), while *A.thiooxidans* prefer acidic environments (pH=3 ± 1). Consequently, the binding capacity of nickel to *T.thiooxidans* NB1-3 in a neutral environment is higher than that in an acidic environment [36]. In an acidic environment, the binding capacity of sodium tungstate with *A.thiooxidans*, is ten times greater than that in a neutral environment [32]. In this study, the surface pH of coupon was only reduced to 7.51 ± 0.10 (Group K), which still corresponds to the initial stage of corrosion. At this point, the predominant microbial species may mainly be NSOB, making nickel more effective at inhibiting H_2_S biological oxidation while sodium tungstate exhibits a lesser effect. Copper oxide is a common metal bacteriostatic agent studied in the field of corrosion control for sewer systems. However, there are no studies comparing the dosages of copper oxide, nickel and sodium tungstate. The results of this study demonstrate that, in a neutral environment, copper oxide requires a higher dosage to exert a more pronounced effect. Moreover, a single type of bacteriostatic agent may only have a specific effect on a particular type of microorganism. Consequently, the use of a single type of bacteriostatic agent alone may not comprehensively address the entire corrosion process. Therefore, in different stages of concrete corrosion, the combined use of multiple types of antibacterial agents may yield superior outcomes.

There are some shortcomings in this study. For instance, no coarse aggregate is added in coupons, which may reduce the cohesion of coupon. The exposure time was not long enough to observe more obvious corrosion differences between different coupons. There was no in-depth analysis of the microbial species in the corrosion layer. In subsequent experiments, larger coupons can be used for longer exposure time to obtain more sufficient experimental data and conclusions. At present, the antibacterial research of most metal bacteriostatic agent is still in the laboratory stage, and the practical application is rare. In order to fully verify the antibacterial effect of metal bacteriostatic agent, field tests are also needed.

## 5. Conclusions

In this study, three metal bacteriostatic agents were added to cement mortar, and then their inhibition effects on MICC were investigated in a laboratory corrosion chamber. The main findings from are:

- The corrosion development process of coupons during exposure generally aligned with the classical three-stage corrosion theory. In the first stage of corrosion development, chemical oxidation of H_2_S primarily drives the process, and the inhibitory effects of different metal bacteriostatic agents on the biological oxidation of H_2_S do not exhibit significant differences at this stage.
- In the second stage, after 28 days of exposure, the surface pH, sulfate concentration, and ATP content of C3 (9.56 ± 0.53, 4.81 mg/cm^2^, and 9.31×10^−6^ ± 0.86×10^−6^ μmol/cm^2^, respectively,) closely resemble those of N1 (9.94 ± 0.02, 6.48 mg/cm^2^, and 10.12×10^−6^ ± 0.78×10^−6^ μmol/cm^2^, respectively,). This indicates that C3 and N1 exhibit superior inhibition of H_2_S biooxidation compared to other coupons of the same type with different dosages. The surface pH, sulfate concentration, and ATP content of W3 are 9.64 ± 0.08, 6.49 mg/cm^2^, and 56.94×10^−6^ ± 5.19×10^−6^ μmol/cm^2^, respectively, which are not superior to those of W1 and W2.
- In the third stage, W3, with the lowest ATP content and the lowest sulfate concentration in Group W, surpasses that in Group C and Group N. This suggests, to some extent, that in the initial stage of corrosion, sodium tungstate exhibits a weaker inhibitory effect on H_2_S biological oxidation compared to copper oxide and nickel oxide.
- Different types of metal bacteriostatic agents have varying effects on H_2_S bio-oxidation at different stages of corrosion development. A single type of bacteriostatic agent may struggle to fully address the entire corrosion process. Consequently, the combined use of multiple bacteriostatic agents may yield sustained inhibitory effects on a range of acid-producing microorganisms throughout the extended corrosion process.

## Acknowledgement

This research is supported by National Key R&D Program of China (2022YFC3203200), and Powerchina key research and development program (DJ-ZDXM-2023-39).

## References

[1]. Dodd, M.C., Potential impacts of disinfection processes on elimination and deactivation of antibiotic resistance genes during water and wastewater treatment. Journal of Environmental Monitoring, 2012. 14(7): p. 1754–1771.

[2]. O Connell, M., C. McNally and M.G. Richardson, Biochemical attack on concrete in wastewater applications: A state of the art review. Cement and Concrete Composites, 2010. 32(7): p. 479–485.

[3]. Breitenmoser, A., et al., Outbreak of acute gastroenteritis due to a washwater-contaminated water supply, Switzerland, 2008. Journal of water and health, 2011. 9(3): p. 569–576.

[4]. Ahmed, W., et al., Quantitative microbial risk assessment of microbial source tracking markers in recreational water contaminated with fresh untreated and secondary treated sewage. Environment international, 2018. 117: p. 243–249.

[5]. Jiang, G., et al., Odor emissions from domestic wastewater: A review. Critical Reviews in Environmental Science and Technology, 2017. 47(17): p. 1581–1611.

[6]. Zhang, L., et al., Chemical and biological technologies for hydrogen sulfide emission control in sewer systems: a review. Water research, 2008. 42(1-2): p. 1–12.

[7]. Liang, S., L. Zhang and F. Jiang, Indirect sulfur reduction via polysulfide contributes to serious odor problem in a sewer receiving nitrate dosage. Water research, 2016. 100: p. 421–428.

[8]. Ling, A.L., et al., Carbon dioxide and hydrogen sulfide associations with regional bacterial diversity patterns in microbially induced concrete corrosion. Environmental science & technology, 2014. 48(13): p. 7357–7364.

[9]. Grengg, C., et al., Advances in concrete materials for sewer systems affected by microbial induced concrete corrosion: A review. Water research, 2018. 134: p. 341–352.

[10]. Wu, M., et al., Microbiologically induced corrosion of concrete in sewer structures: A review of the mechanisms and phenomena. Construction and Building Materials, 2020. 239: p. 117813.

[11]. Ling, A.L., et al., High-resolution microbial community succession of microbially induced concrete corrosion in working sanitary manholes. PloS one, 2015. 10(3): p. e0116400.

[12]. Okabe, S., et al., Succession of sulfur-oxidizing bacteria in the microbial community on corroding concrete in sewer systems. Applied and environmental microbiology, 2007. 73(3): p. 971–980.

[13]. Roberts, D.J., et al., Quantifying microbially induced deterioration of concrete: initial studies. International Biodeterioration & Biodegradation, 2002. 49(4): p. 227–234.

[14]. Valix, M., et al., Microbiologically induced corrosion of concrete and protective coatings in gravity sewers. Chinese Journal of Chemical Engineering, 2012. 20(3): p. 433–438.

[15]. Diercks, M., W. Sand and E. Bock, Microbial corrosion of concrete. Experientia, 1991. 47: p. 514–516.

[16]. Yu, L., et al., The differential proliferation of AOB and NOB during natural nitrifier cultivation and acclimation with raw sewage as seed sludge. RSC advances, 2020. 10(47): p. 28277–28286.

[17]. Valix, M., Analysis of acid transport through multi-phase epoxy mortars for wastewater structures. Water Science and Technology, 2015. 72(2): p. 332–337.

[18]. Gutiérrez-Padilla, M.G.D., et al., Biogenic sulfuric acid attack on different types of commercially produced concrete sewer pipes. Cement and Concrete Research, 2010. 40(2): p. 293–301.

[19]. De Muynck, W., N. De Belie and W. Verstraete, Effectiveness of admixtures, surface treatments and antimicrobial compounds against biogenic sulfuric acid corrosion of concrete. Cement and Concrete Composites, 2009. 31(3): p. 163–170.

[20]. Gevaudan, J.P., et al., Copper and cobalt improve the acid resistance of alkali-activated cements. Cement and Concrete Research, 2019. 115: p. 327–338.

[21]. Volland, S., et al., Intracellular chromium localization and cell physiological response in the unicellular alga Micrasterias. Aquatic toxicology, 2012. 109: p. 59–69.

[22]. Cloete, T.E., Resistance mechanisms of bacteria to antimicrobial compounds. International Biodeterioration & Biodegradation, 2003. 51(4): p. 277–282.

[23]. Vittoriadiamanti, M. and M.P. Pedeferri, Concrete, mortar and plaster using titanium dioxide nanoparticles: applications in pollution control, self-cleaning and photo sterilization, in Nanotechnology in eco-efficient construction. 2013, Elsevier. p. 299–326.

[24]. Sikora, P., et al., Antimicrobial activity of Al2O3, CuO, Fe3O4, and ZnO nanoparticles in scope of their further application in cement-based building materials. Nanomaterials, 2018. 8(4): p. 212.

[25]. Singh, V.P., et al., Photocatalytic, hydrophobic and antimicrobial characteristics of ZnO nano needle embedded cement composites. Construction and Building Materials, 2018. 158: p. 285–294.

[26]. Haile, T., et al., Evaluation of the bactericidal characteristics of nano-copper oxide or functionalized zeolite coating for bio-corrosion control in concrete sewer pipes. Corrosion Science, 2010. 52(1): p. 45–53.

[27]. Vishwakarma, V., et al., Enhancing antimicrobial properties of fly ash mortars specimens through nanophase modification. Materials Today: Proceedings, 2016. 3(6): p. 1389–1397.

[28]. Noeiaghaei, T., et al., Biogenic deterioration of concrete and its mitigation technologies. Construction and Building Materials, 2017. 149: p. 575–586.

[29]. Qiu, L., et al., Antimicrobial concrete for smart and durable infrastructures: A review. Construction and Building Materials, 2020. 260: p. 120456.

[30]. Alum, A., et al., Cement-based biocide coatings for controlling algal growth in water distribution canals. Cement and Concrete Composites, 2008. 30(9): p. 839–847.

[31]. Delgado, K., et al., Polypropylene with embedded copper metal or copper oxide nanoparticles as a novel plastic antimicrobial agent. Letters in applied microbiology, 2011. 53(1): p. 50–54.

[32]. Negishi, A., et al., Growth inhibition by tungsten in the sulfur-oxidizing bacterium Acidithiobacillus thiooxidans. Bioscience, biotechnology, and biochemistry, 2005. 69(11): p. 2073–2080.

[33]. Sugio, T., et al., Mechanism of growth inhibition by tungsten in Acidithiobacillus ferrooxidans. Bioscience, biotechnology, and biochemistry, 2001. 65(3): p. 555–562.

[34]. Gyu-Yong, K., et al., Evaluation of properties of concrete using fluosilicate salts and metal (Ni, W) compounds. Transactions of Nonferrous Metals Society of China, 2009. 19: p. s134–s142.

[35]. Kong, L., B. Zhang and J. Fang, Effect of bactericide on the deterioration of concrete against sewage. Journal of Materials in Civil Engineering, 2018. 30(8): p. 04018160.

[36]. Maeda, T., et al., Nickel inhibition of the growth of a sulfur-oxidizing bacterium isolated from corroded concrete. Bioscience, biotechnology, and biochemistry, 1996. 60(4): p. 626–629.

[37]. Kong, L., B. Zhang and J. Fang, Study on the applicability of bactericides to prevent concrete microbial corrosion. Construction and Building Materials, 2017. 149: p. 1–8.

[38]. Dehbozorgi, M. and R. Bazargan Lari, Investigating the effect of using colors containing copper oxide and zinc oxide (zinc-rich) on microbial corrosion behavior of sewage pipes. journal of New Materials, 2019. 10(35): p. 15–26.

[39]. Li, X., et al., The rapid chemically induced corrosion of concrete sewers at high H2S concentration. Water research, 2019. 162: p. 95–104.

[40]. Sun, X., et al., Effects of surface washing on the mitigation of concrete corrosion under sewer conditions. Cement and Concrete Composites, 2016. 68: p. 88–95.

[41]. Cayford, B.I., et al., Comparison of microbial communities across sections of a corroding sewer pipe and the effects of wastewater flooding. Biofouling, 2017. 33(9): p. 780–792.

[42]. Jiang, G., J. Keller and P.L. Bond, Determining the long-term effects of H2S concentration, relative humidity and air temperature on concrete sewer corrosion. Water Research, 2014. 65: p. 157–169.

[43]. Islander, R.L., et al., Microbial ecology of crown corrosion in sewers. Journal of Environmental Engineering, 1991. 117(6): p. 751–770.

[44]. Joseph, A.P., et al., Surface neutralization and H2S oxidation at early stages of sewer corrosion: Influence of temperature, relative humidity and H2S concentration. Water Research, 2012. 46(13): p. 4235–4245.

[45]. Wells, T. and R.E. Melchers, An observation-based model for corrosion of concrete sewers under aggressive conditions. Cement and Concrete Research, 2014. 61–62: p. 1-10.

[46]. Nogami, Y., et al., Inhibition of sulfur oxidizing activity by nickel ion in Thiobacillus thiooxidans NB1-3 isolated from the corroded concrete. Bioscience, biotechnology, and biochemistry, 1997. 61(8): p. 1373–1375.

[47]. Jacob, F., The Role of Sulfur Oxidizing Bacteria on Corrosion of X65 Low Carbon Steels and its Mitigation Using Sodium Tungstate and Nickel Biocides. 2013.

