## Supplementary information for "Comparative study on the inhibition of copper oxide, nickel, and sodium tungstate on microbially induced concrete corrosion under sewer conditions"

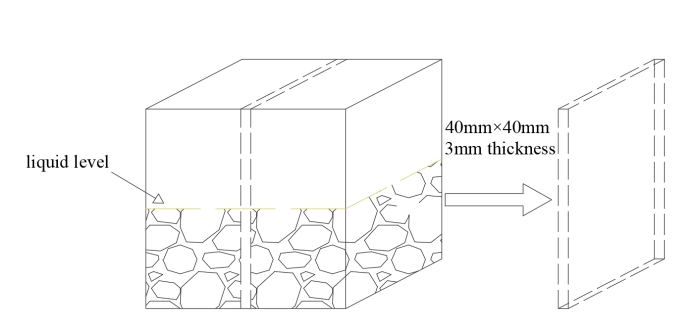

Fig. S1 The concrete samples were used as a slicing mode for SEM and EDS detection.

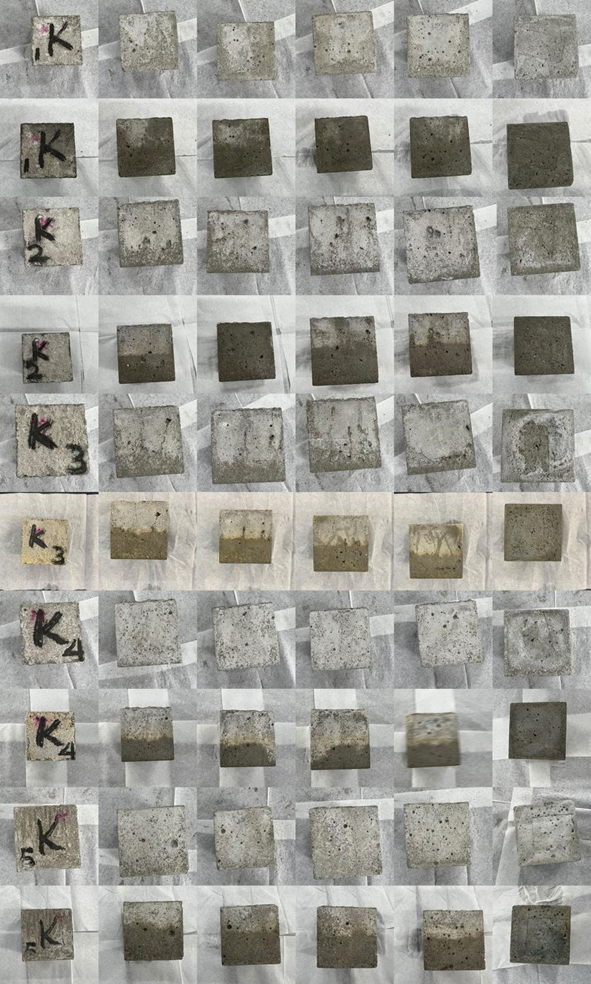

(A)

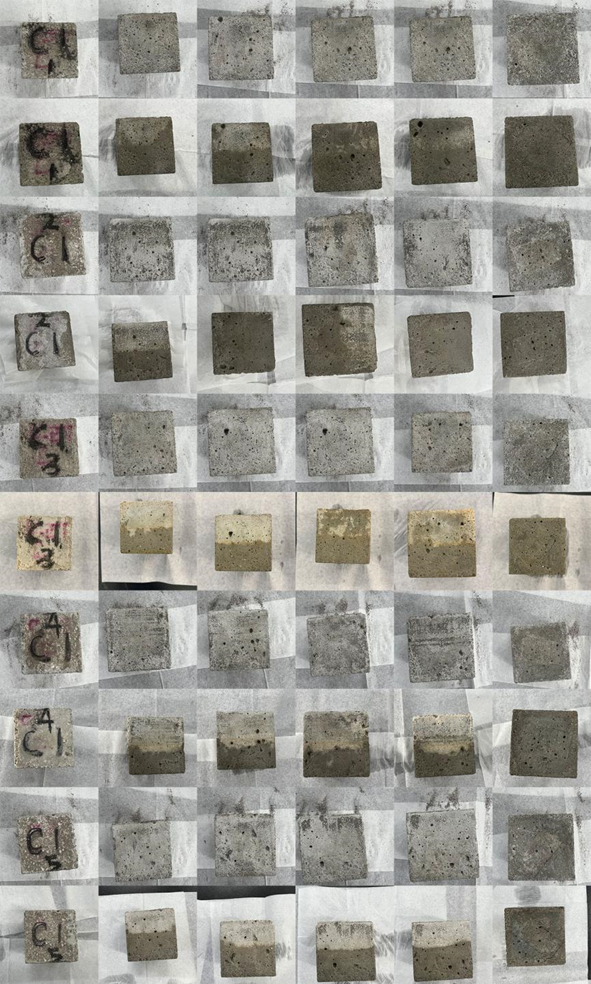

(B)

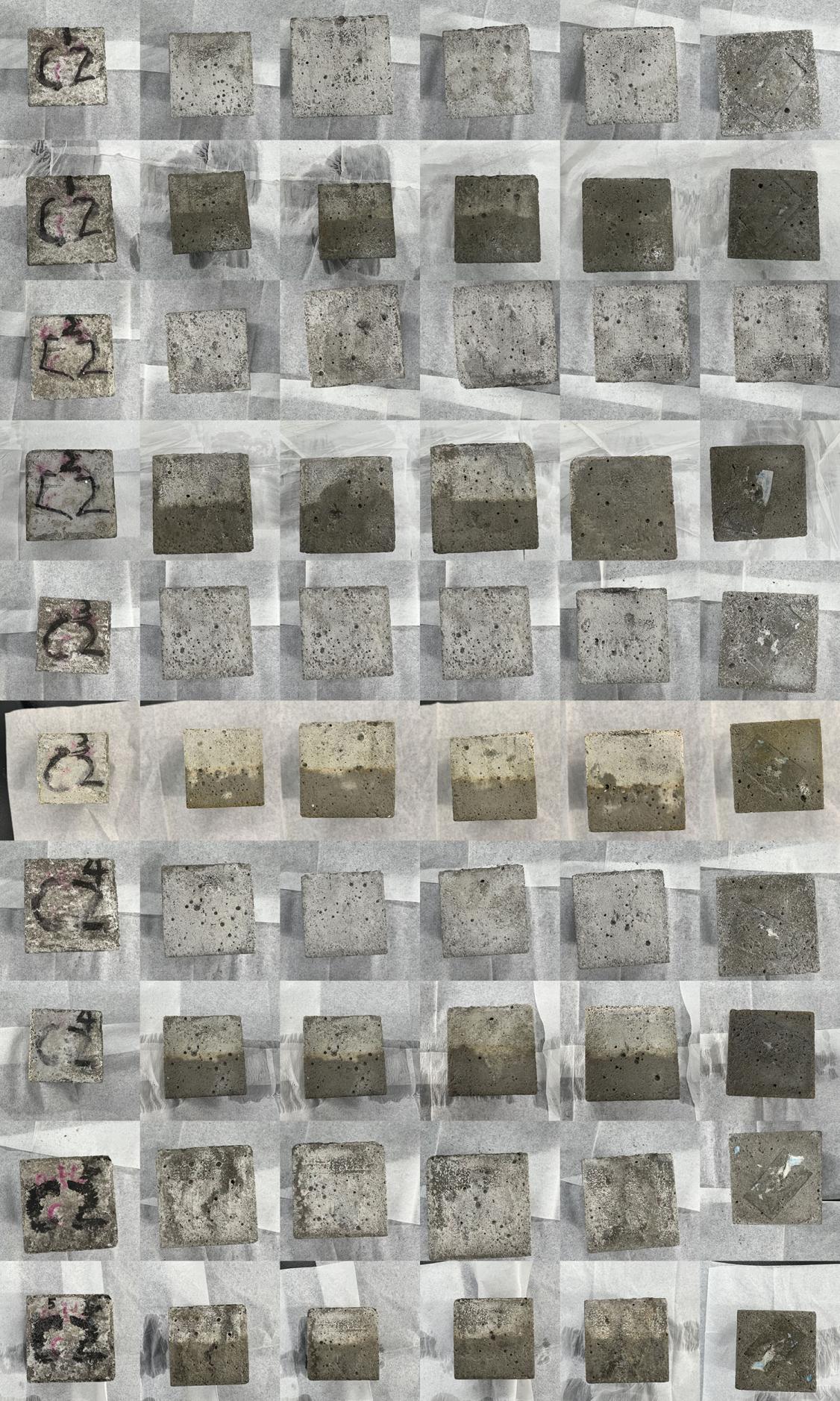

(C)

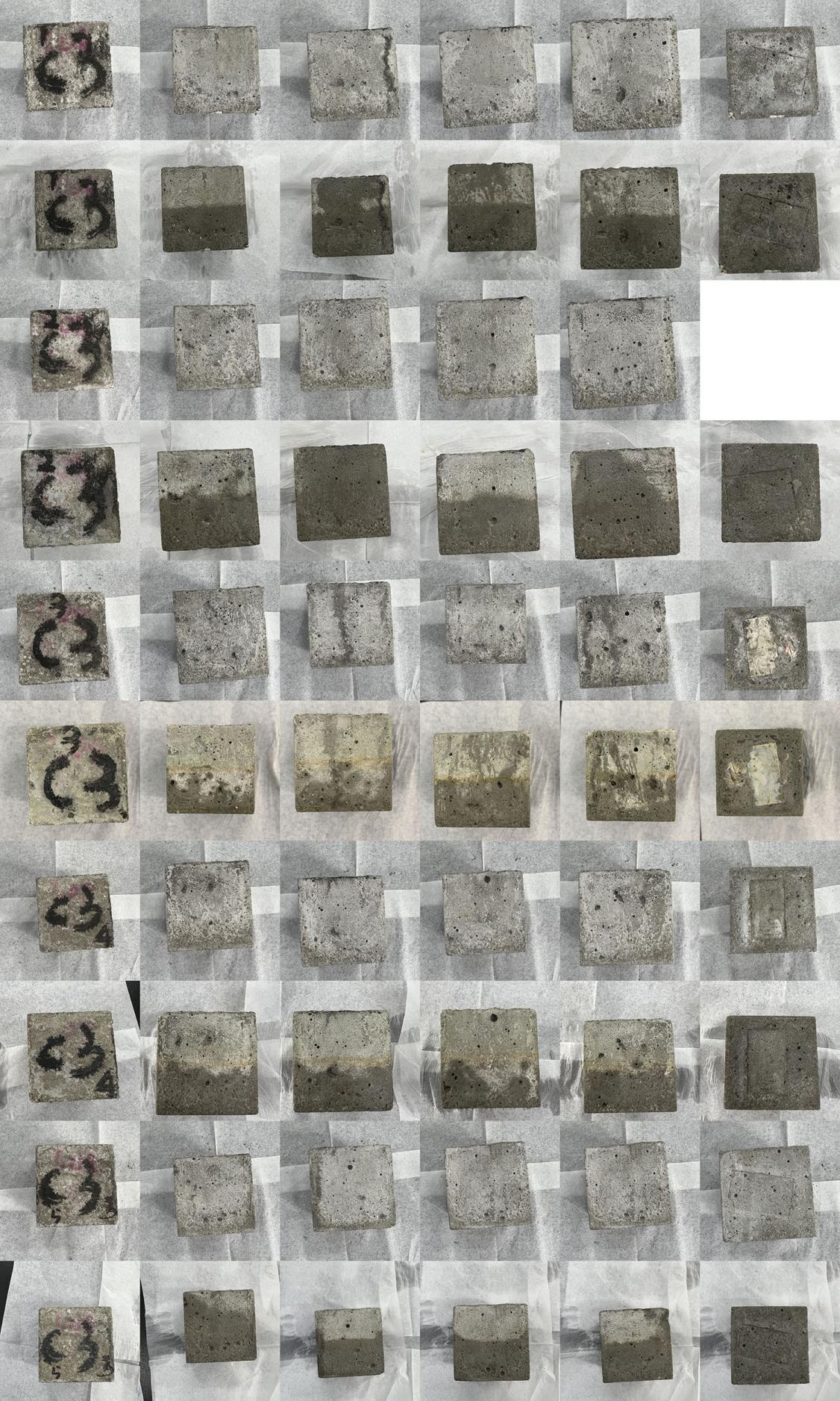

(D)

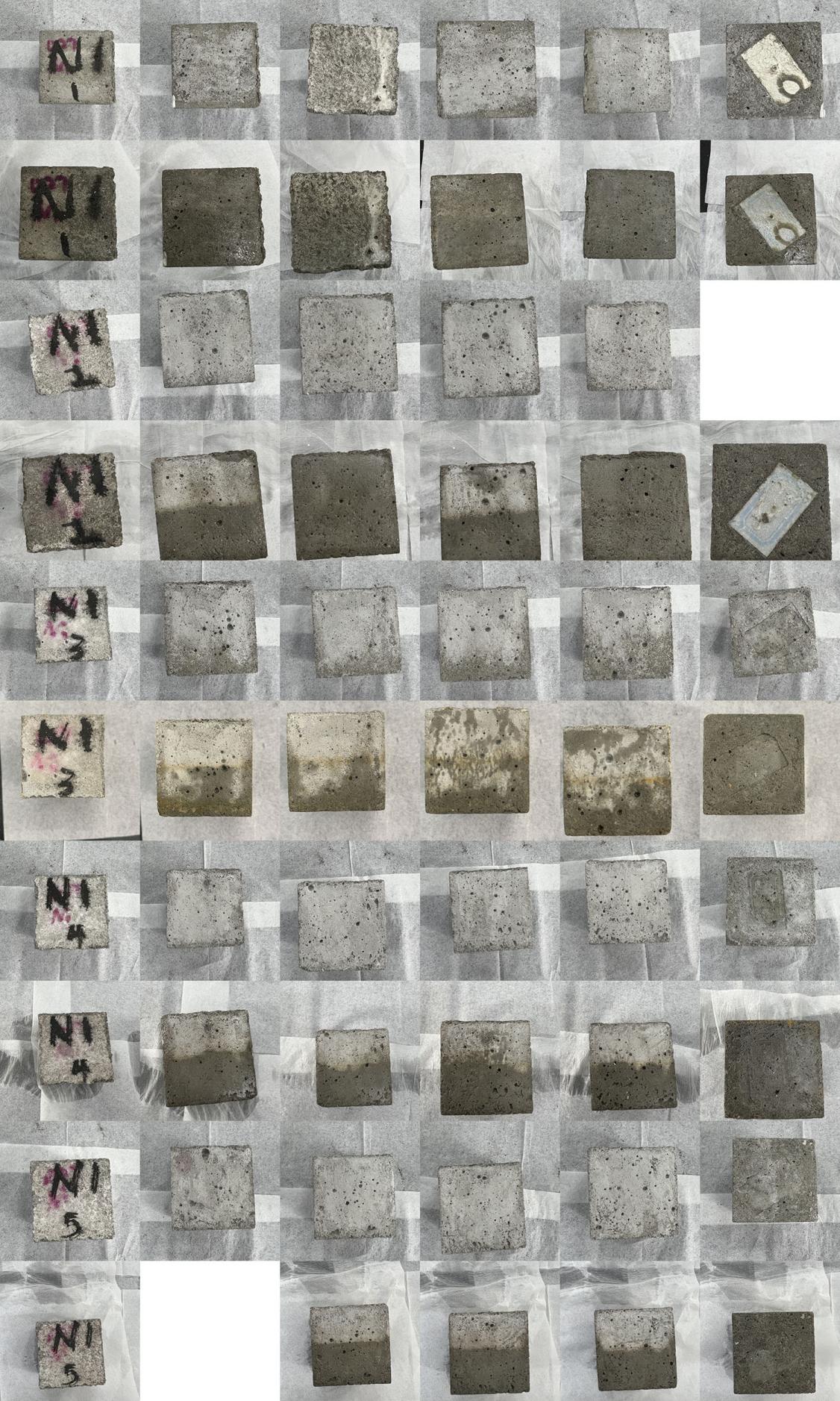

(E)

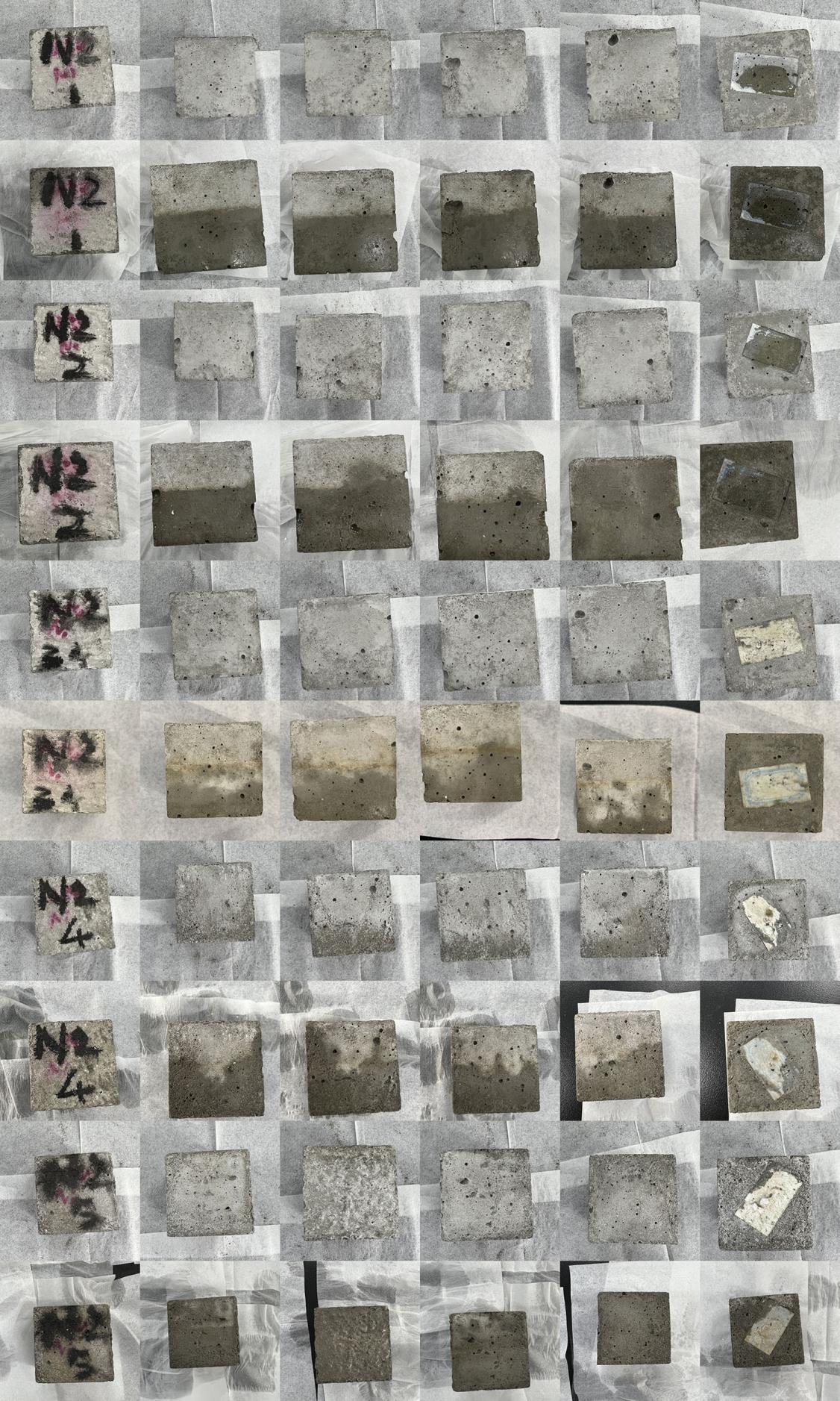

(F)

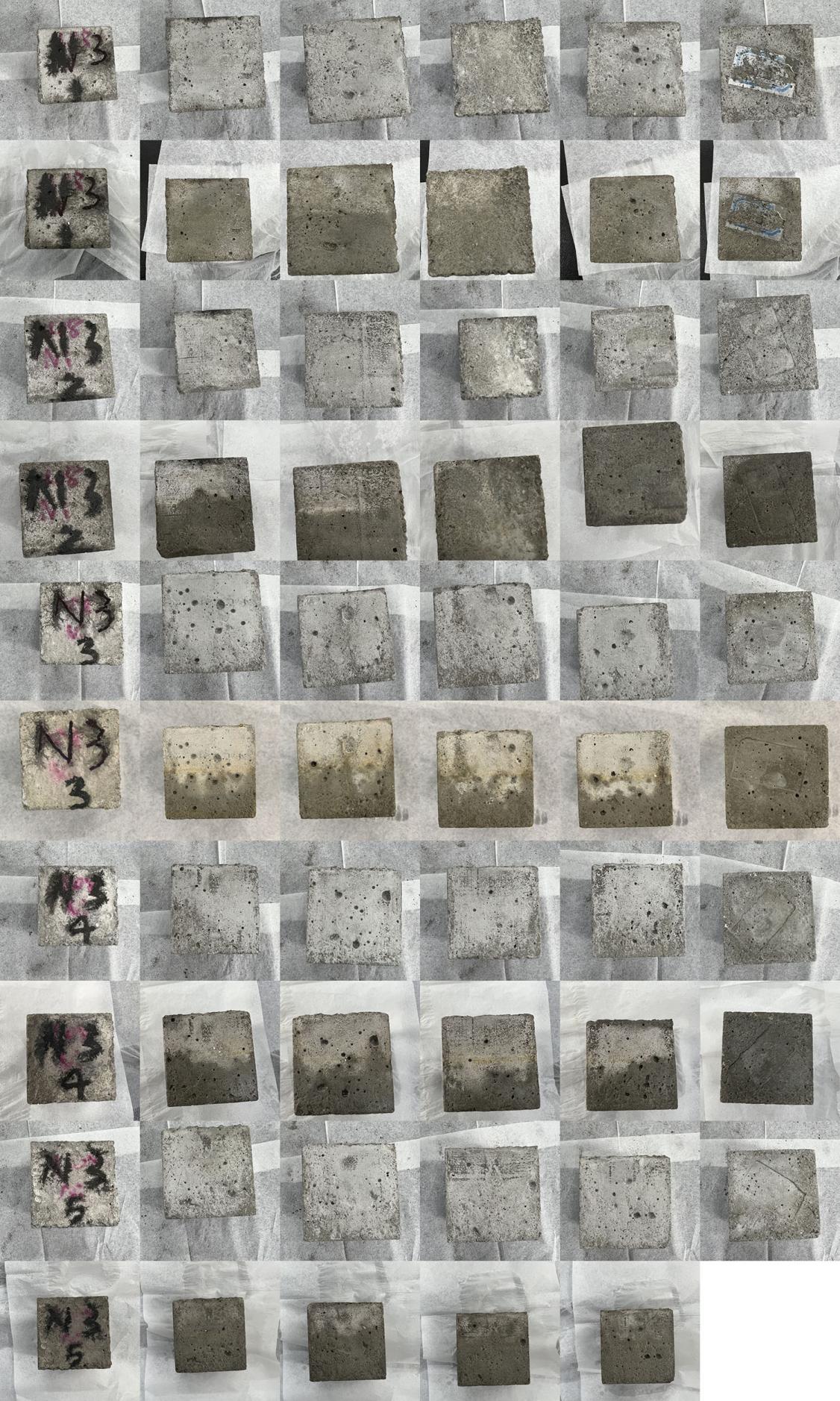

(G)

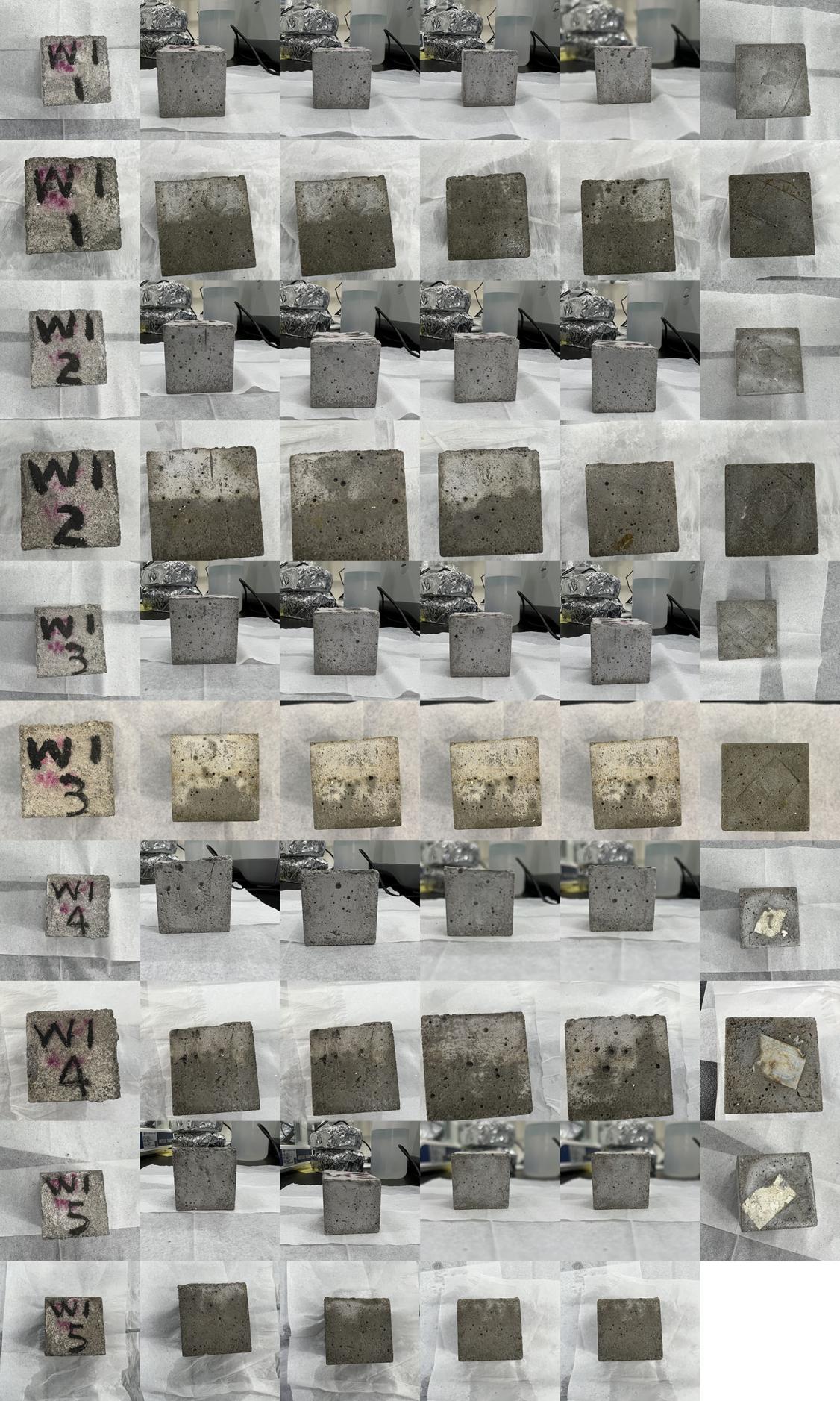

(H)

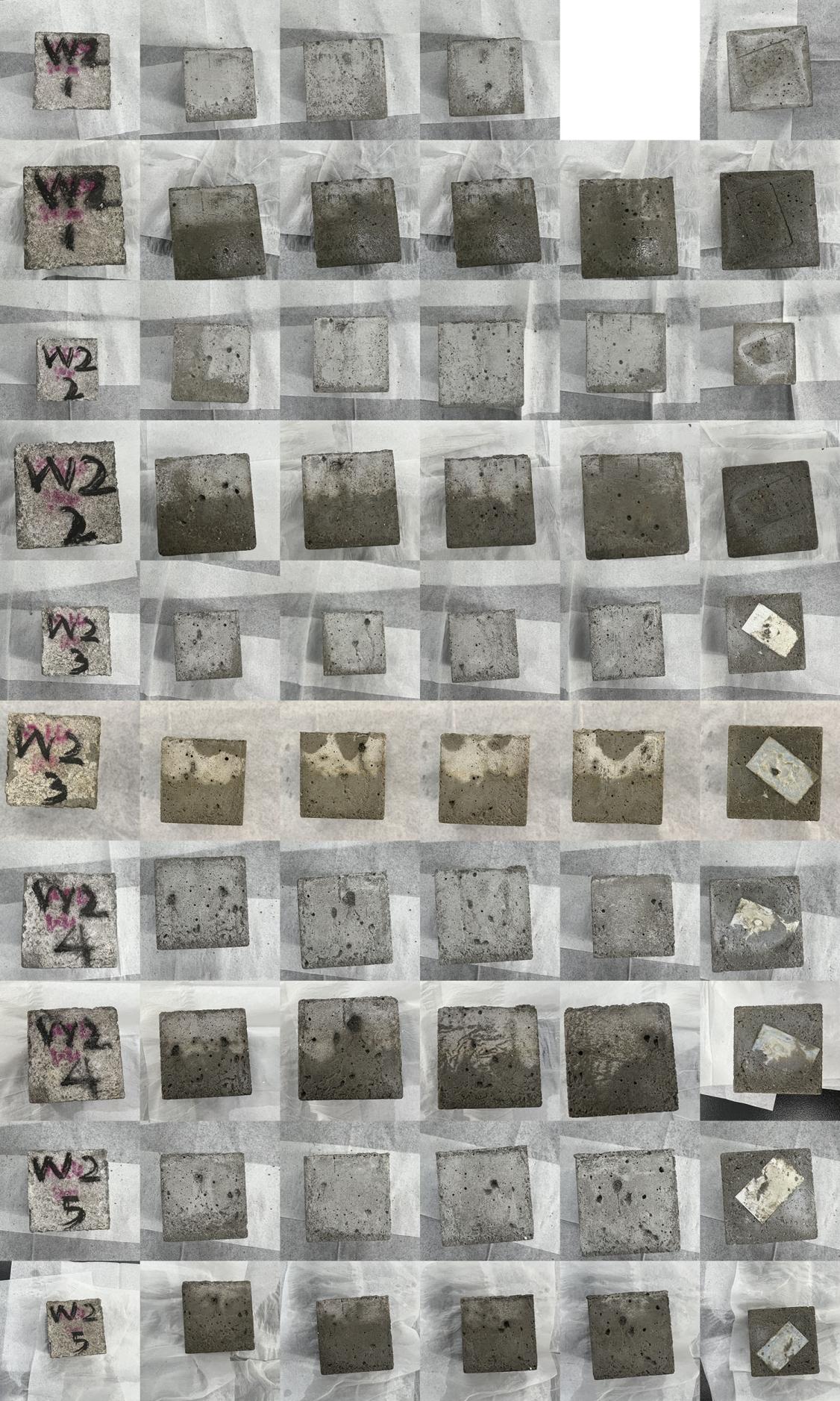

(I)

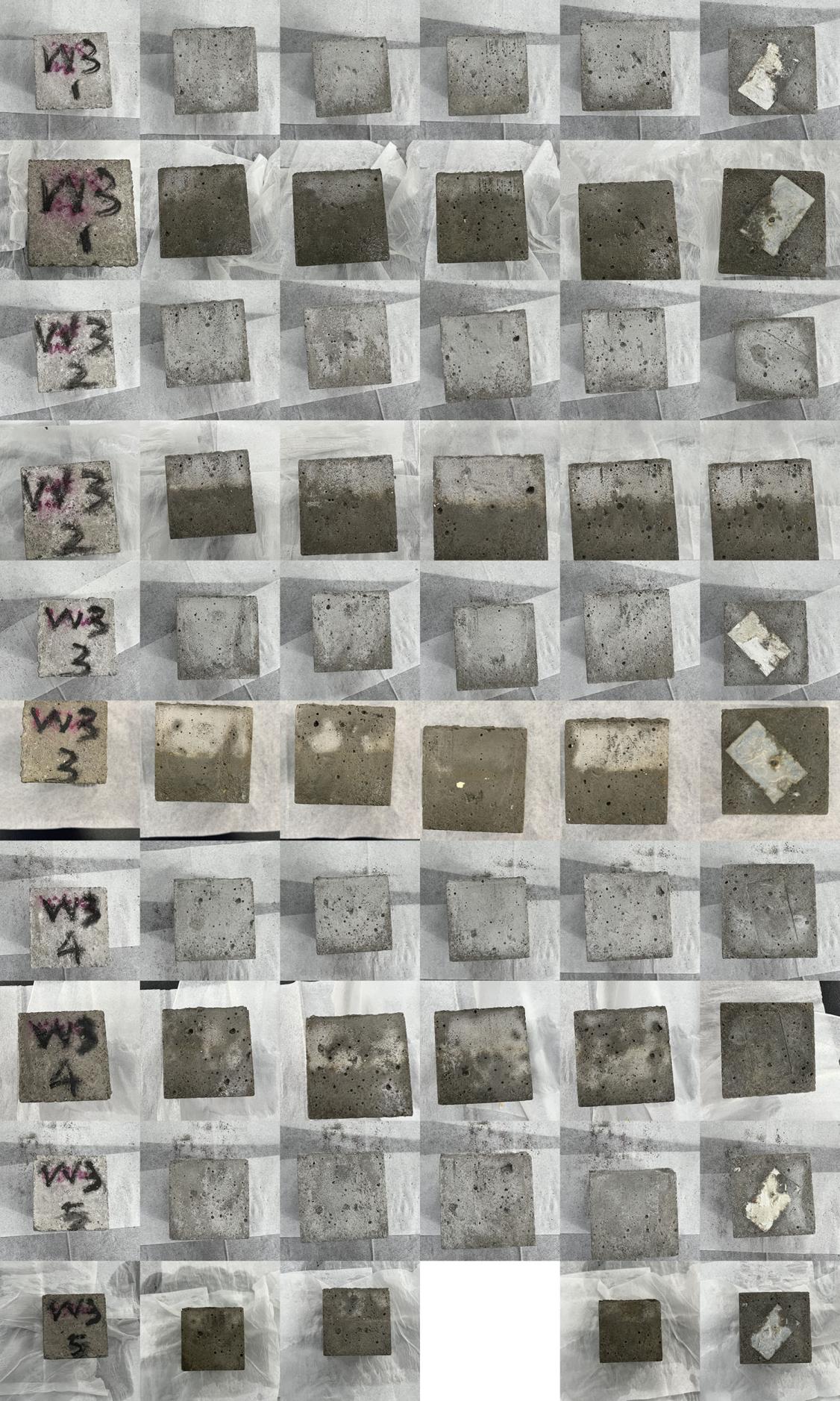

(J)

Fig. S2 The initial state and appearance change of the same sample during the corrosion process. The number 1/2/3/4/5 on the sample indicates that the sample has experienced 7/14/28/42/56 days of corrosion. (A) control group K, (B) experimental group C1 (add 0.05% copper oxide), (C) experimental group C2 (add 0.1% copper oxide), (D) experimental group C3 (add 0.2% copper oxide), (E) experimental group N1 (add 0.05% nickel), (F) experimental group N2 (add 0.1% nickel), (G) experimental group N3 (add 0.2% nickel), (H) experimental group W1 (add 0.05% sodium tungstate), (I) experimental group W2 (add 0.1% sodium tungstate), (J) experimental group W3 (add 0.2% sodium tungstate).

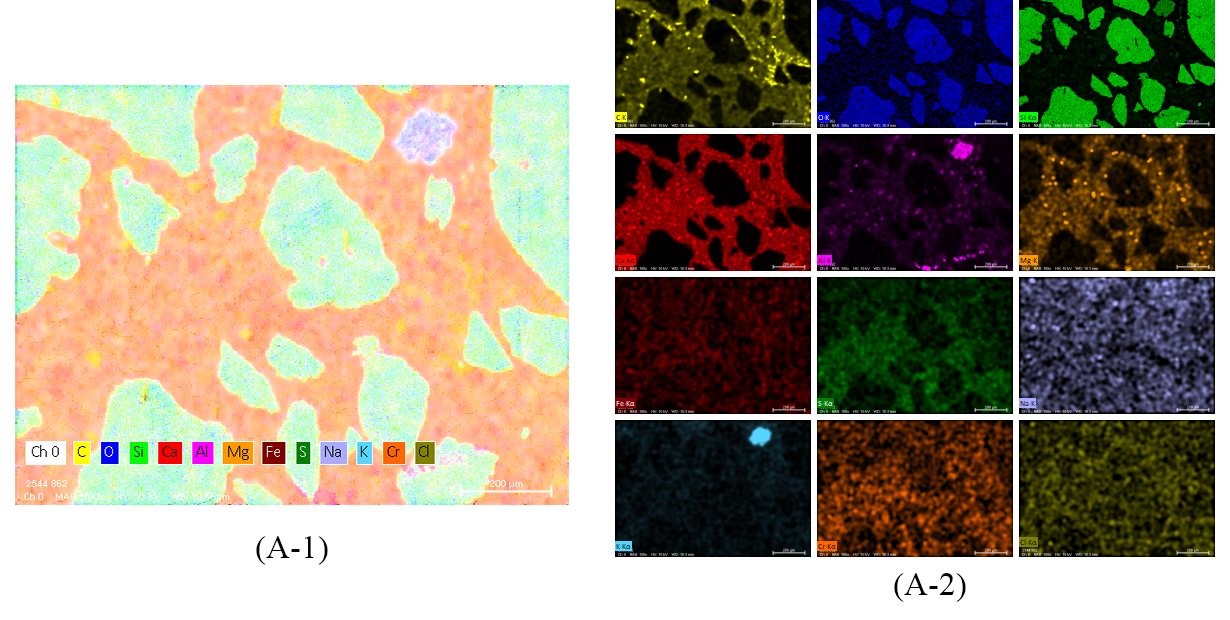

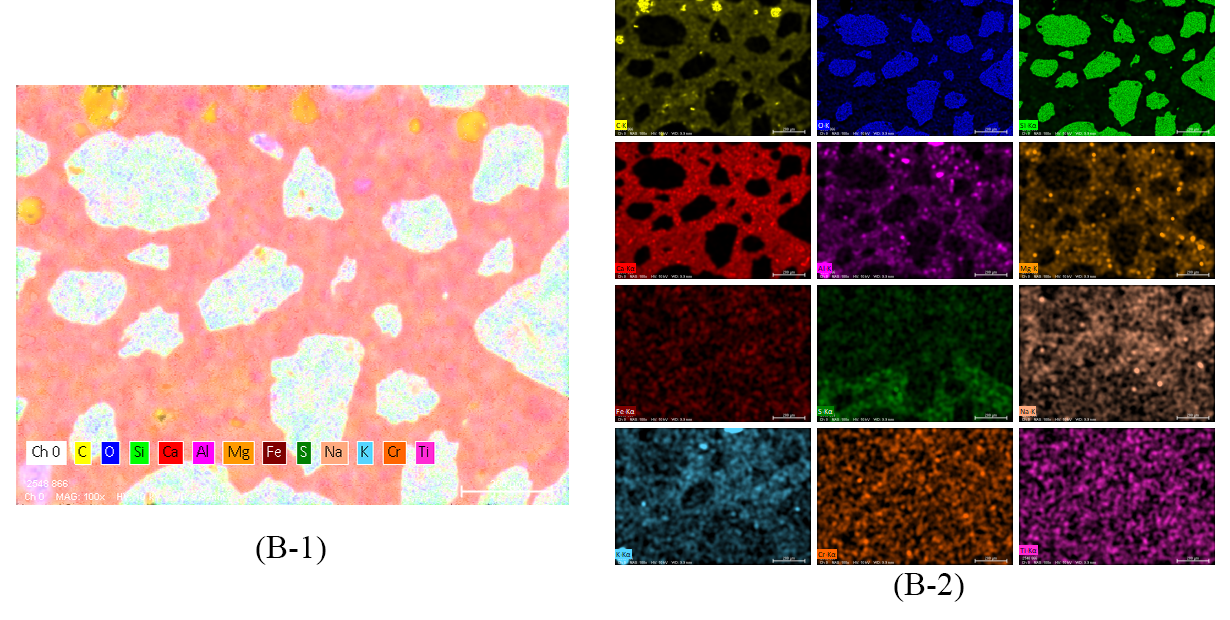

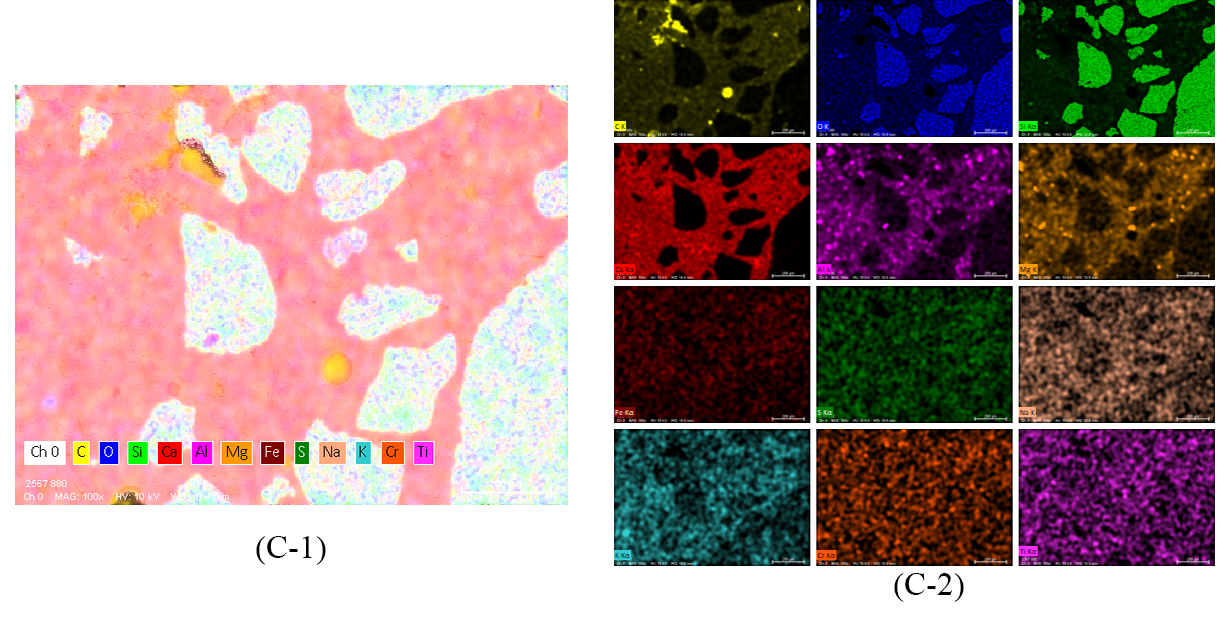

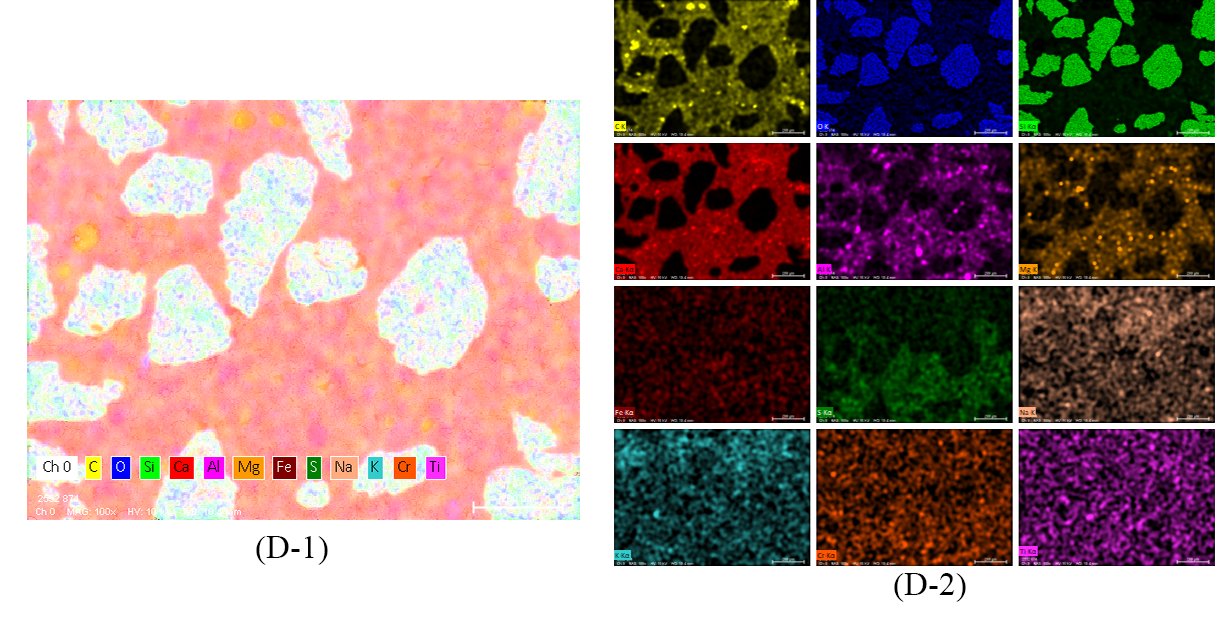

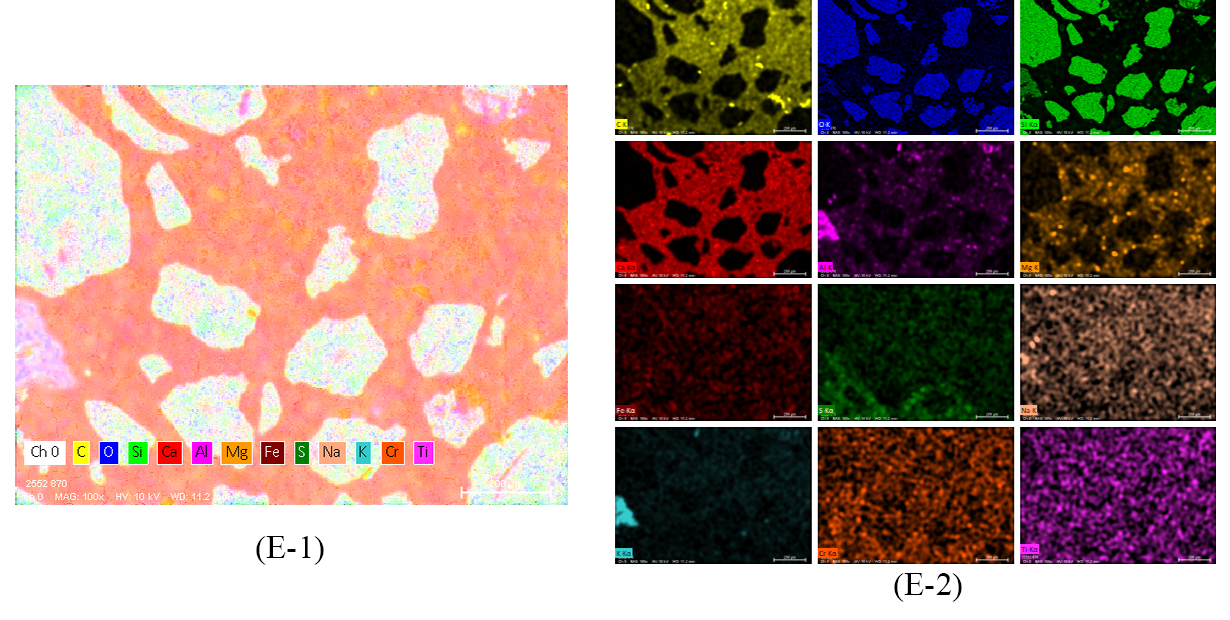

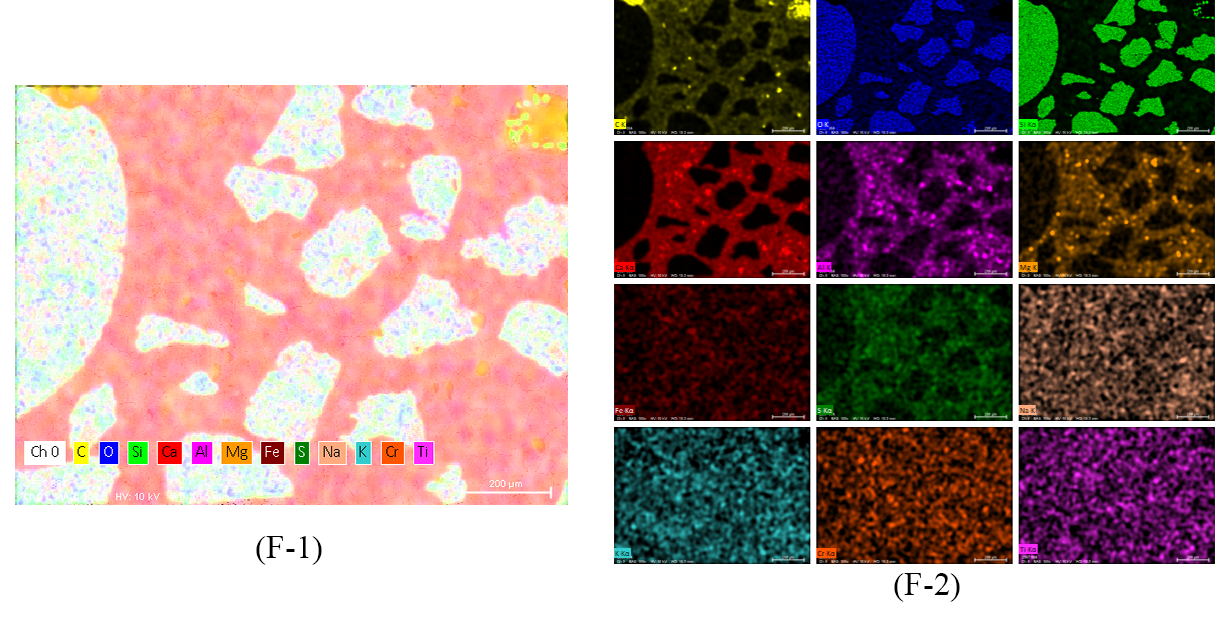

Fig. S3 Energy spectrum diagram of the region by the SEM in Fig.8 and the number 1 is the composite energy spectrum of all elements, and the number 2 is the individual energy spectrum of each element.

Table S1. Physical indexes of the bactericide.

| Bactericide | Molecular formula | Appearance | Density (g/cm^3^) | Solubility | Cost  (Yuan/100g)* |
| --- | --- | --- | --- | --- | --- |
| Copper oxide | CuO | Black powder | 6.31 | Insoluble | 169.60 |
| Nickel | Ni | Black or gray-black powder | 8.90 | Insoluble | 65.60 |
| Sodium tungstate | NaWO_4_ | White crystal or powder | 3.25 | Soluble | 63.20 |

*The price is the price of high-purity chemicals purchased for this study and is for reference only.

Table S2. The indexes of domestic sewage (mg / L).

| Sampling time | COD | NH_3_ - N | Chloride | Temperature  (℃) |
| --- | --- | --- | --- | --- |
| 2022.01.05 | 128 | 18.41 | 372.8 | 21 |
| 2022.01.26 | 158 | 18.01 | 136.3 | 20 |
| 2022.03.09 | 93 | 19.01 | 106.5 | 20 |
| 2022.03.31 | 86 | 13.97 | 102.2 | 22 |

Table S3. The normalized mass of various elements on EDS (%).

| The normalized mass | K0^a^ | K56^b^ | C3 | N1 | N3 | W3 |
| --- | --- | --- | --- | --- | --- | --- |
| C | 12.41 | 16.78 | 12.08 | 15.58 | 11.86 | 13.95 |
| O | 46.74 | 45.45 | 47.74 | 45.25 | 48.77 | 45.15 |
| Na | 0.27 | 0.44 | 0.50 | 0.39 | 0.35 | 0.30 |
| Mg | 0.58 | 0.66 | 0.74 | 0.74 | 0.67 | 0.77 |
| Al | 0.96 | 0.93 | 0.95 | 0.96 | 1.03 | 0.92 |
| Si | 23.41 | 20.30 | 21.31 | 21.49 | 21.98 | 24.06 |
| S | 0.23 | 0.29 | 0.12 | 0.53 | 0.19 | 0.50 |
| Cl | 0.01 | - | - | - | - | - |
| K | 0.24 | 0.30 | 0.28 | 0.14 | 0.28 | 0.11 |
| Ca | 14.00 | 13.98 | 15.50 | 13.95 | 13.97 | 13.26 |
| Ti | - | 0.05 | 0.04 | 0.02 | 0.07 | 0.06 |
| Cr | 0.49 | 0.21 | 0.24 | 0.30 | 0.20 | 0.27 |
| Fe | 0.67 | 0.60 | 0.51 | 0.65 | 0.62 | 0.64 |
| Total | 100 | 100 | 100 | 100 | 100 | 100 |

^a^ K0 is the blank control group before exposure; ^b^K56 is the blank control group after 56 days of exposure.
